# *Caenorhabditis elegans* locomotion is affected by internalized paramagnetic nanoparticles in the presence of magnetic field

**DOI:** 10.1101/248369

**Authors:** Eleni Gourgou, Yang Zhang, Ehsan Mirzakhalili, Bogdan Epureanu

## Abstract

*C. elegans* nematodes are a model organism used broadly to investigate the impact of environmental factors on physiology and behavior. Here, *C. elegans* with internalized paramagnetic nanoparticles were placed inside magnetic field to explore effects on locomotion. We hypothesize that internalized paramagnetic nanoparticles combined with external magnetic field affect *C. elegans*’ locomotion machinery. To test our hypothesis, we used young adult *C. elegans* fed on bacteria mixed with paramagnetic nanoparticles of 1 μm, 100 nm and 40 nm diameter. The presence of nanoparticles inside the worms’ body (alimentary canal, body muscle) was verified by fluorescent and electron microscopy. A custom-made software was used to track freely moving *C. elegans* in the absence or presence of magnetic field sequentially for 200+200 sec. We used established metrics to quantify locomotion-related parameters, including posture, motion and path features. Key features of *C. elegans* locomotion (increased body bends and stay ratio, decreased range, forward movement, and speed along the magnetic field) were affected in worms with internalized nanoparticles of 100 nm and 1 μm in the presence of magnetic field, in contrast to untreated worms. Our work contributes on clarifying the effect of internalized paramagnetic nanoparticles, combined with magnetic field, on *C. elegans* locomotion.

**Summary Statement:** *C. elegans* with internalized paramagnetic nanoparticles are placed inside magnetic field to explore effects on locomotion. Results support the potential of *C. elegans* to investigate the impact of the above environmental factors on behavior.

## Introduction

The effects of magnetic field (MF) on living organisms have been a target of numerous research efforts, with their number increasing significantly during the last decades (Ghodbane et al., 2013; Hong, 1995; Shaw et al., 2015). In addition to the interest scientific community shows on the effect of alternating MF on cells (Ueno et al., 1986; Öcal, 2008; Belova and Acosta-Avalos, 2015), static MF effects have gained attention also (Miyakoshi, 2005; Teodori et al., 2002) mainly due to their correlation with activity linked to the modern way of living (Lewczuk, 2014). The type of MF least studied in regard to cells is high-gradient MF. Recent work provides the theoretical framework for the possible impact of high-gradient MF of various sources on cells’ molecular components and function (Zablotskii et al., 2016).

Model organisms have been a successful resource to study MF effects on various types of cells and tissues (Osipova et al., 2016; Shcherbakov et al.; Malkemper et al., 2015; Kumari et al., 2017). Invertebrate models, like *Drosophila melanogaster*, have been used since the 80’s (Fedele et al., 2014; Naito et al., 2012; Ramirez et al., 1983; Kale and Baum, 1980; Bae et al., 2016; Giachello et al., 2016). Interestingly, even though the nematode *C. elegans* has been an emblematic model organism to study the impact of a plethora of stimuli and environmental factors on behavior and physiology (Cheung et al., 2005; De Bono and Bargmann, 1998; Liedtke et al., 2003; Hedgecock and Russell, 1975; Ward et al., 2008), only recently it has been used in MF related work (Vidal-Gadea, 2015; Njus, 2015; Long et al., 2015; Wang et al., 2015; Lee et al., 2010), in which the first animal magnetosensory neurons were identified (Vidal-Gadea, 2015). The presence of biogenic magnetite has also been reported in *C. elegans* (Cranfield et al., 2004).

Nanoparticles uptake by *C. elegans* worms has been a successful means to evaluate toxicity of heavy metals and pollutants (Khare et al., 2011; Meyer et al., 2010; Kim et al., 2017), and the importance of *C. elegans* as a model system for *in vivo* nanoparticle assessment has been specifically highlighted (Gonzalez-Moragas et al., 2015a). Worms’ behavior (Ma et al., 2009) and locomotion (Li et al., 2012; Wu et al., 2012) have been evaluated under the influence of internalized metal nanoparticles. In addition, magnetized nanoparticles have been used to activate ion channels in *C. elegans* through heating (Huang et al., 2010). However, only very recently internalized nanoparticles were used to locally enhance MF in the worms’ body and study the subsequent impact on its metabolism (Wang et al., 2017).

*C. elegans* locomotion has been a major behavioral output used to investigate the impact of genetic background, environmental factors and diverse treatments on the worm’s nervous system (Gourgou, 2016; Li et al., 2016; Liu, 2013; Hsu et al., 2009; Parida et al., 2014; Pierce-Shimomura et al., 2008). Locomotion features have been characterized and quantified extensively and are being used as an indicator of *C. elegans* physiological status and healthspan (Bansal et al., 2015; Shtonda and Avery, 2006; Peliti et al., 2013; Hahm et al., 2015). Therefore, locomotion is one of the first behaviors investigated to define whether *C. elegans* nematodes are sensitive to an environmental factor of interest.

The abovementioned scientific premises, the remaining need to clarify MF effects on animal physiology, and the increasing interest in the sensitivity of *C. elegans* to MF, indicate that the investigation of MF gradient effects on worms’ behavior comes at a mature point. We hypothesize that internalized paramagnetic nanoparticles combined with external magnetic field affect *C. elegans*’ locomotive behavior. To test our hypothesis, we used internalized paramagnetic nanoparticles that can generate an effect inside the worms’ body, in the closest possible proximity with tissues and cells. We used locomotion as a quantifiable and revealing behavioral expression to determine the effect of MF combined with the internalized nanoparticles. Our results demonstrate a response of *C. elegans* locomotion machinery to internalized paramagnetic nanoparticles in combination with MF gradients, and they pave the way for future studies seeking to clarify the participation of excitable cells, muscles and potentially even neurons, to this still uncharacterized behavior.

## Results

### Magnetic field gradient characterization

The simulation results for both electromagnets agreed with the experimental results provided by the manufacturer (Fig. S1). An overview of the MF around the electromagnets and the geometry of the model in COMSOL are presented in Figure 2A and a clear view of the MF on the plane of the worm plate surface is presented in Figure 2B. The arrows demonstrate the direction of the MF between the two electromagnets. The contours show that the MF is stronger near the electromagnets, as expected.

**Fig. 1:**
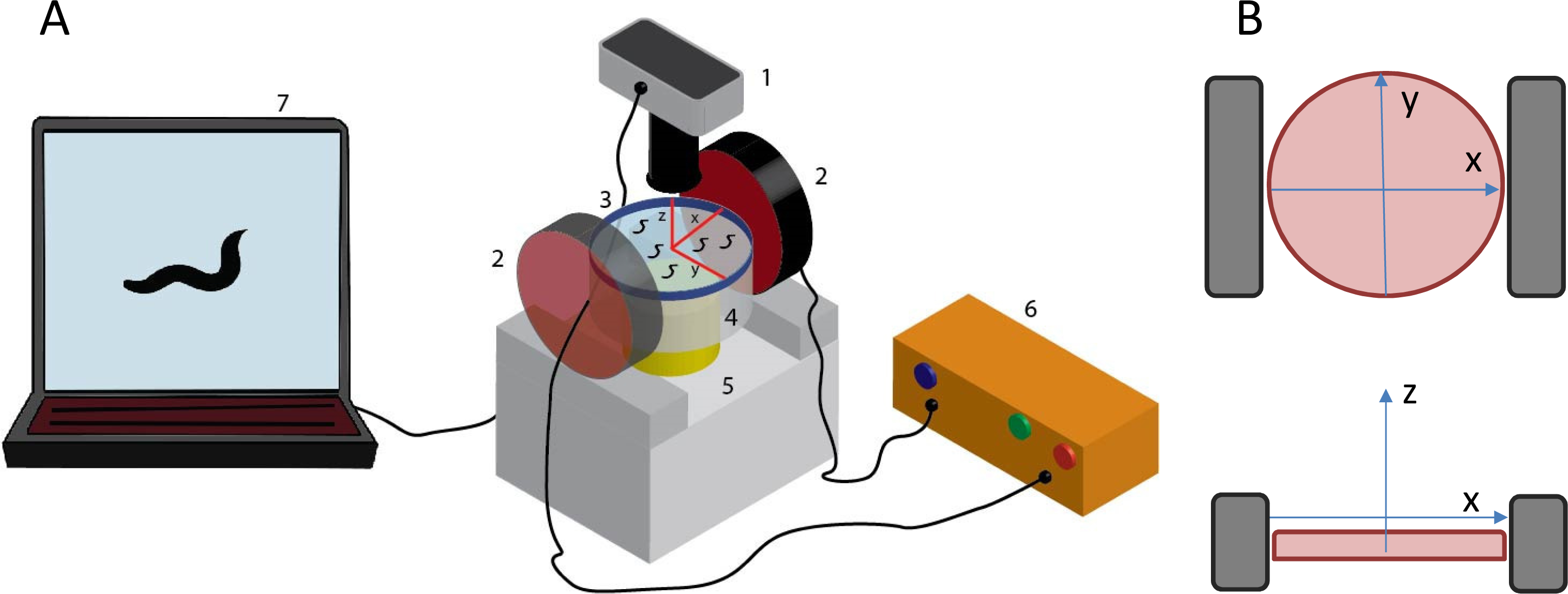
Experimental set up. A: Parts of the experimental set up for the application of gradient magnetic field on freely moving *C. elegans.* 1: Objective lens and camera; 2: Electromagnets; 3: NGM plate with freely moving wild type N2 *C. elegans*, with schematic of plate orientation, red lines indicating x, y, and z axes; 4: Auxiliary transparent base; 5: Working stage with bright light source; 6: Power Supply; 7: Computer and recording software. Objects are not depicted in scale. B: Schematic of the orientation of the NGM plate (pink circle), x, y, and z axes and electromagnets (grey rectangles), top: view from above, bottom: view from aside.

**Fig. 2:**
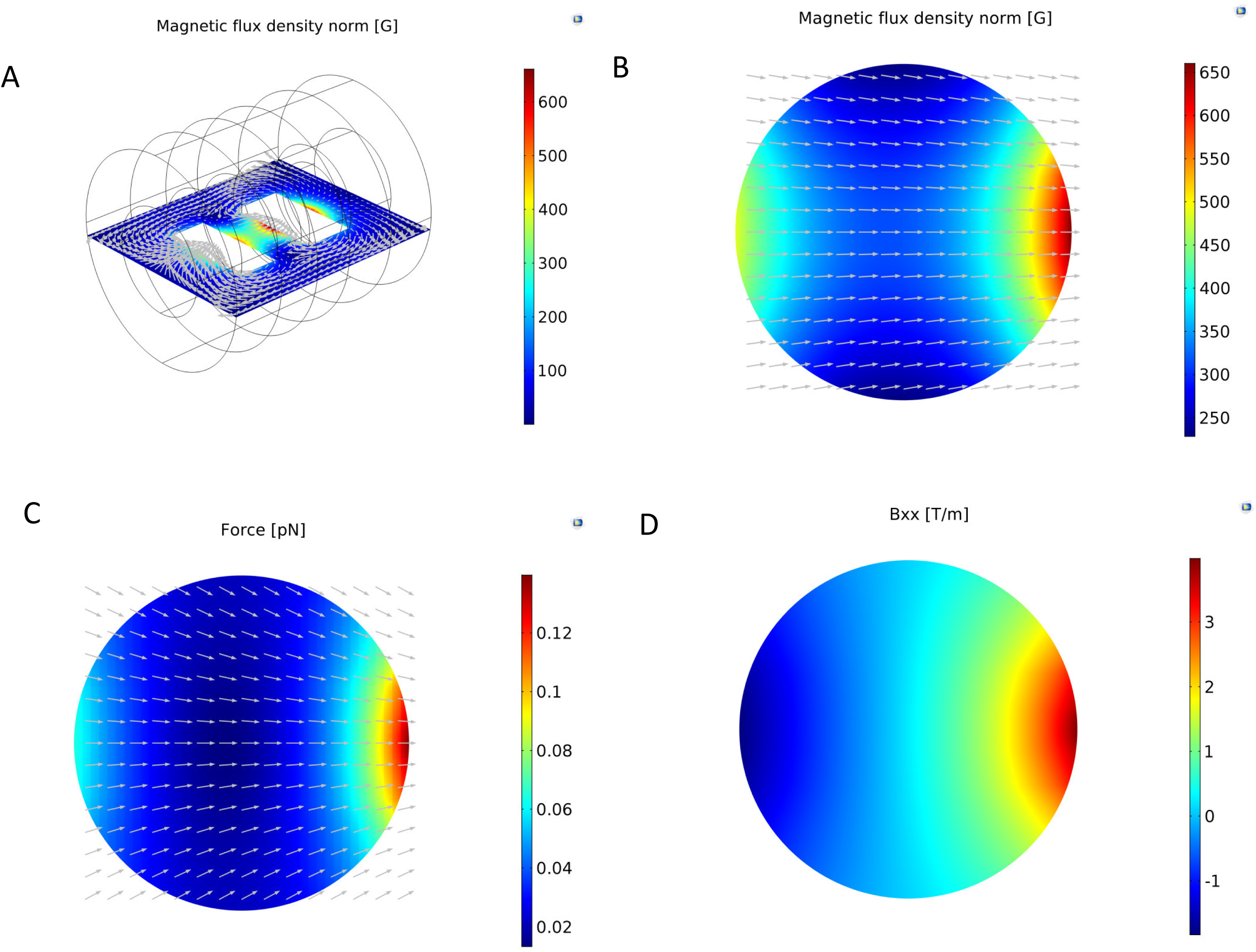
COMSOL Multiphysics simulation results for the magnetic field generated by the electromagnets. A: Overview of the magnetic field flux density on the plane of the worm plate surface. The arrows show the direction of the magnetic flux. B: The magnetic field flux density distribution on the worm plate surface. The arrows indicate the direction of the magnetic field (the component of the magnetic field in the perpendicular direction is set equal to zero to avoid arrows going in/out of the plane). C: The magnetic forces applied on particles located on the plane of the worm plate surface. The arrows show the direction of the magnetic forces (the component of the force in the perpendicular direction is set equal to zero to avoid arrows going in/out of the plane). D: The gradient of the magnetic field in the direction of the axis that connects the centers of the two electromagnets.

We focused on the features of the MF and the forces generated on the plate surface, where the worms’ locomotion takes place. The MF was almost one-dimensional on the plate surface and was stronger nearer the electromagnets (Fig. 2C). There were 9 components for the gradient of the MF. In Figure 2D, the strongest component of the MF gradient is shown (Bxx), which is parallel to *x* axis. The magnitude of the gradient was larger near the electromagnets.

The nanoparticles create secondary MFs in the presence of an external MF. Details on calculating the forces that were created by the particles used in the present study can be found in the Supplementary Information section. The magnitude of the MF flux was calculated in MATLAB using Eqs. [2] and [3] of Supplementary Information for configurations along the *x* and *y* axes of three nanoparticles (as shown in Fig. 3A–3D). The MF was stronger close to the particles for both configurations and decayed rapidly as the distance from the particles increased (Fig. 3E). The force between the particles in the *x* direction was attractive, while the force between the particles in the *y* direction was repulsive. The attractive forces between the particles allow them to form chain-like structures, if they are not interrupted by the medium in which the particles are located (Mirzakhalili et al., 2017; Nakata et al., 2008). The magnetic moment of the external MF for the particles on the worm plate surface is depicted in Figure 3F.

**Fig. 3:**
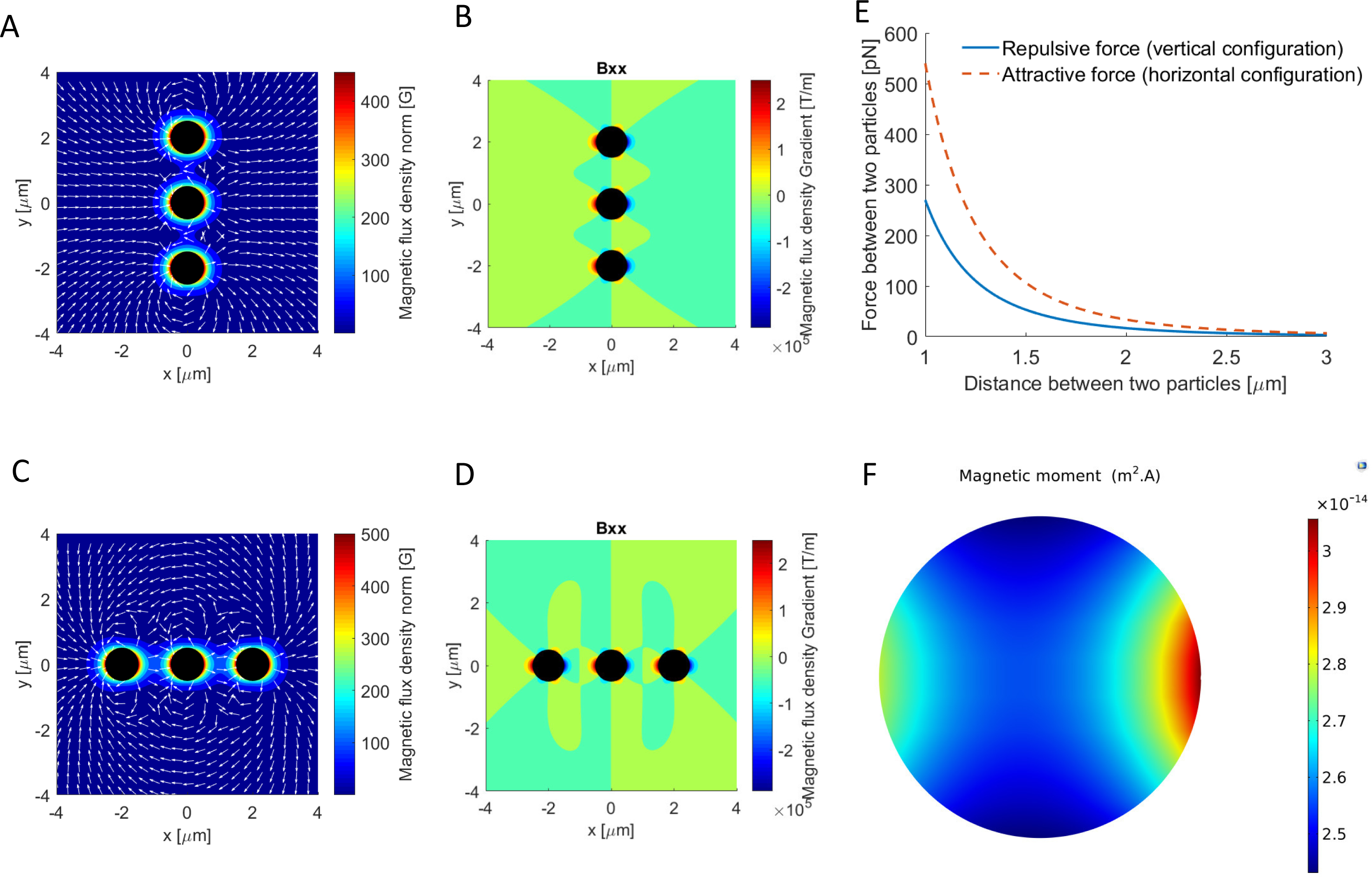
Characterization of the magnetic field around the 1μm nanoparticles for two different configurations. The direction of magnetic moment for both configurations is along the x axis, as is shown in Figure 1. The magnetic moment of the particles is assumed to be similar and equal to the maximum value that is computed from the COMSOL Multiphysics simulations in the plate. A: The magnetic field flux density around three paramagnetic particles in the vertical configuration, i.e. along the y axis. The arrows indicate the direction of the magnetic field. B: The largest component of the gradient of the magnetic field for the vertical configuration of the paramagnetic particles. C: The magnetic field flux density norm around three paramagnetic particles in the horizontal configuration, i.e. along the x axis. The arrows indicate the direction of the magnetic field. D: The largest component of the gradient of the magnetic field for the horizontal configuration of the paramagnetic particles. E: The forces between two particles in each configuration. F: The magnetic moment of the external magnetic field, which the particles experience once inside the magnetic field.

### Confirmation of nanoparticles uptake and particle localization in *C. elegans*’ body

Nanoparticles mixed with bacterial food were successfully internalized, as verified by microscopy methods, selected according to the properties of each particle group (Fig. 4). The presence of 1 μm paramagnetic particles (Table 1) in the worm’s intestine and in the pharynx around the grinder area was verified by bright field microscopy. The particles appear as dark (copper-colored) objects accumulated in the alimentary canal (Fig. 4A, right panel), whereas control animals’ intestine area appears transparent (Fig. 4A, left panel). Uptake of 100 nm fluorescent, paramagnetic particles (Table 1) was verified by fluorescent microscopy. The particles appear to accumulate along the intestine lumen and in the pharynx, as shown when filters for rhodamine, the fluorescent substance with which the particles were coated (see Table 1 for particles properties), were used (Fig. 4B). Successful feeding on 40 nm paramagnetic particles (Table 1) was confirmed by scanning electron microscopy. As shown in Figure 4C, when using the circular backscatter (CBS) detector, 40 nm particles were visualized as white dots under the worm cuticle, in the broad area downstream of the pharynx and along the alimentary canal. The deformation of the sample due to the process followed allowed for obtaining only the approximate location of the particles. The white dots which represent the particles appear in different sizes, which might be attributed to particle aggregates or to the different depth at which the particles were located.

**Fig. 4:**
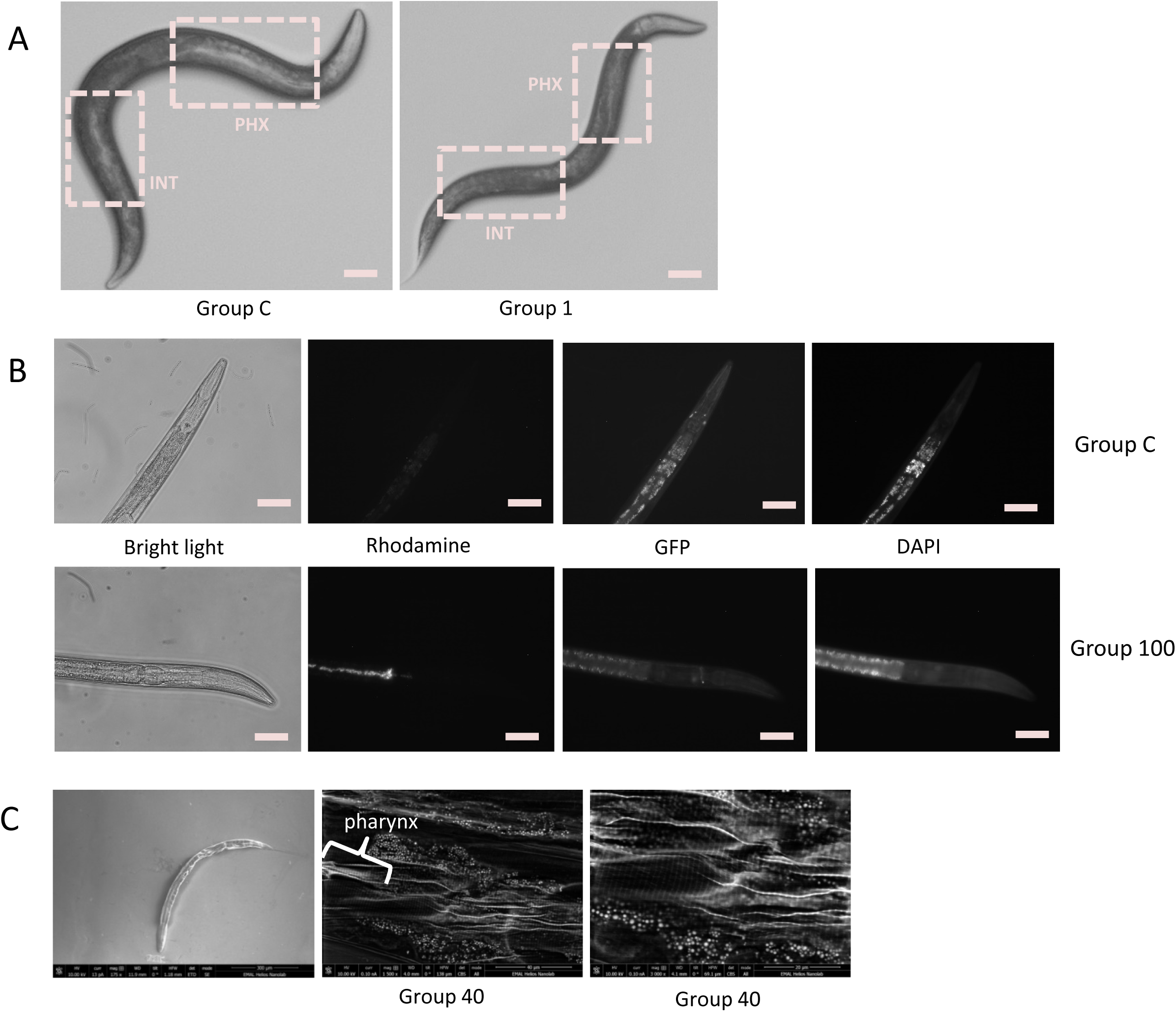
Confirmation of nanoparticles uptake in young adult *C. elegans*. A: Internalization of 1 μm paramagnetic particles is verified by bright field microscopy. Left: worm fed with plain *E. coli* OP50 (control, Group C). Right: worm fed with *E. coli* OP50 mixed with 1um particles. Particles appear to be aggregated in the dark-colored pharynx (PHX) and intestine (INT) of Group 1 worms, in contrast to the light-colored pharynx (PHX) and intestine (INT) of Group C worms. Scale bar: 0.1mm. B: Internalization of 100 nm magnetic, fluorescent nanoparticles is verified by epifluorescent microscopy. Top panels: worm fed with plain *E. coli* OP50 (control, Group C), bottom panels: worm fed with E. coli OP50 mixed with 100 nm particles. Bright light: worms illuminated by bright light source; rhodamine: worms visualized with optical filter for rhodamine, Excitation 545 nm/Emission 565 nm; GFP: worms visualized with optical filter for green fluorescent protein (GFP), Excitation 395 nm/Emission 510 nm; DAPI: worms visualized with optical filter for DAPI, Excitation 358 nm/Emission 460 nm. In GFP and DAPI images, autofluorescence is the only fluorescence detected. Scale bar: 0.1 mm. C: Internalization of 40 nm paramagnetic particles is verified by scanning electron microscopy (SEM). Left: a whole *C. elegans* as captured by SEM, using Everhart-Thornley SE detector. Center: 40 nm particles, shown as white dots, detected close to *C. elegans* pharynx, using circular backscatter (CBS) detector, magnification 1500x. Right: 40 nm particles, shown as white dots, detected close to *C. elegans* pharynx, using circular backscatter (CBS) detector, magnification 3000x. Location of particles is approximate, due to distortion generated during sample processing.

**Table 1:** Groups of worms tested and properties of the respective nanoparticles.

| Group | Particle Size | Coating | Magnetic properties | Fluorescence |
| --- | --- | --- | --- | --- |
| Group C | - | - | - | - |
| Group 1 | 1 µm | Streptavidin | Paramagnetic | No |
| Group 100 | 100 nm | No | Paramagnetic | Rhodamine |
| Group 40 | 40 nm | -COOH | Paramagnetic | No |

To investigate the particles localization in the worms’ body, and to decipher whether they pass the intestine barrier, we used transmission electron microscopy. TEM images show that the particles can be found in the intestine (Fig. 5D) and the intestine lumen (Fig. 5E). Interestingly, particles aggregates were also detected inside muscle tissue, very close to the body wall (Fig. 5F). Therefore, the nanoparticles can be located very close to excitable cells.

**Fig. 5:**
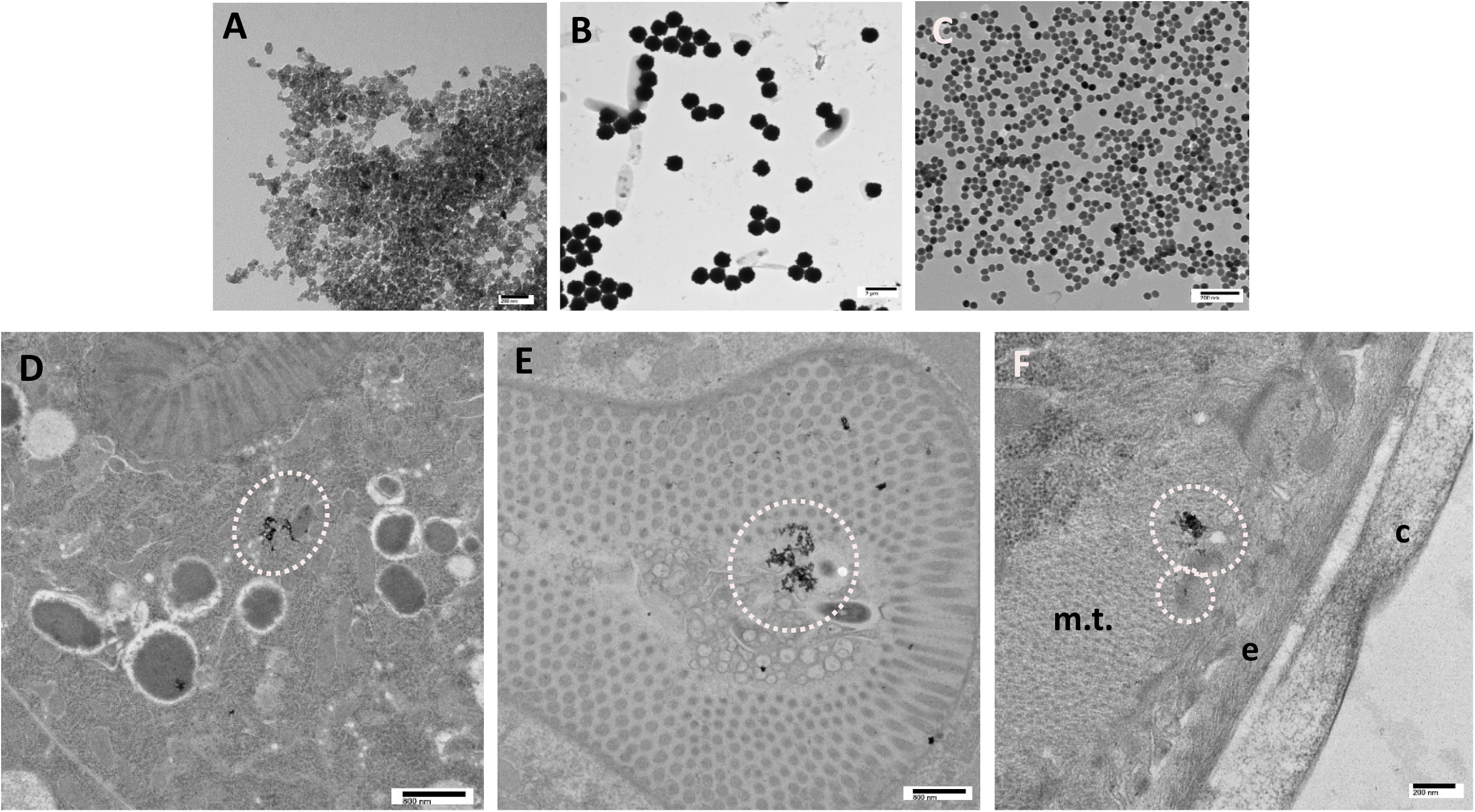
Location of nanoparticles in young adult *C. elegans*, using Transmission Electron Microscopy (TEM). A-B-C: Free particles (A: 100 nm, B: 1 μm, C: 40 nm) imaged with TEM; D: 40 nm particles aggregates in the intestine; E: 100 nm particles aggregates in the intestine lumen; F: 40 nm particles aggregate in body muscle tissue (m.t.), close to epidermis (e) and cuticle (c). Dotted circles indicate particles aggregates in all panels.

### Analysis of *C. elegans* locomotion

Analysis of selected locomotion features revealed that worms fed with 1 μm and 100 nm diameter paramagnetic nanoparticles, when they moved freely in MF, had altered locomotion dynamics, compared to worms without internalized nanoparticles.

We examined selected posture features for each worm of each group, namely the total body bends in degrees, and the number of bends (bend count) realized per worm. The total body bends were not affected by the presence of either particles or of MF (Fig. 6A). However, there was a significant increase in the number of bends per worm of Group 100 (Fig. 6B, WSR test *p*-value = 0.031) when the MF was on.

**Fig. 6:**
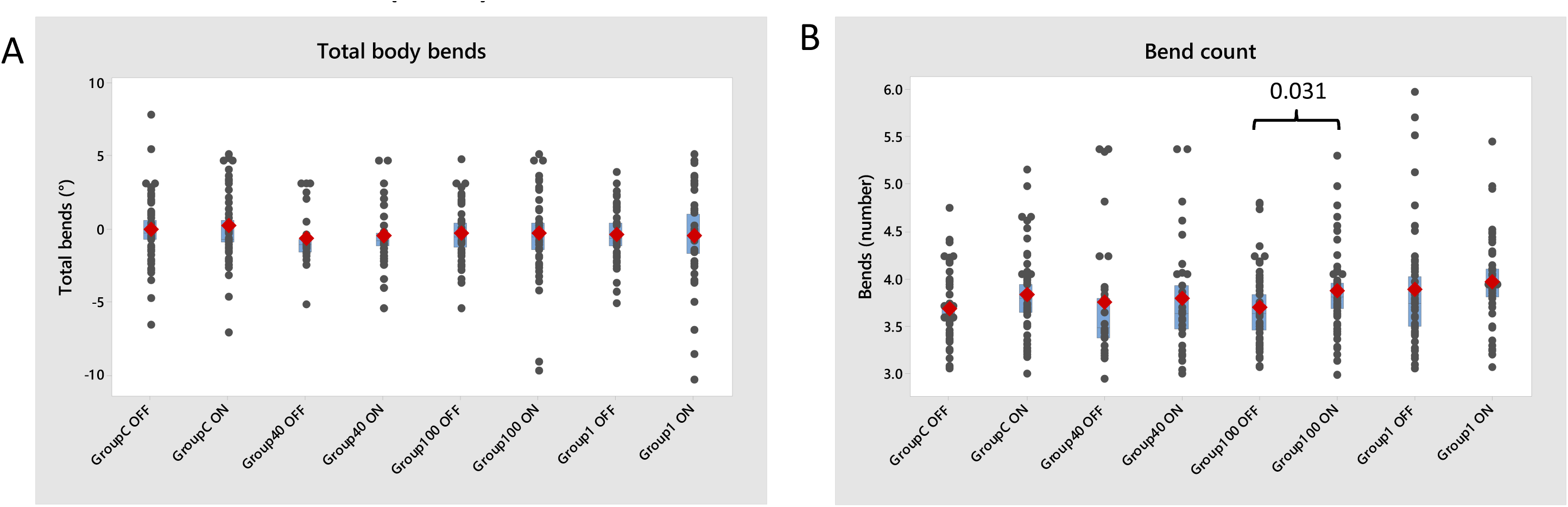
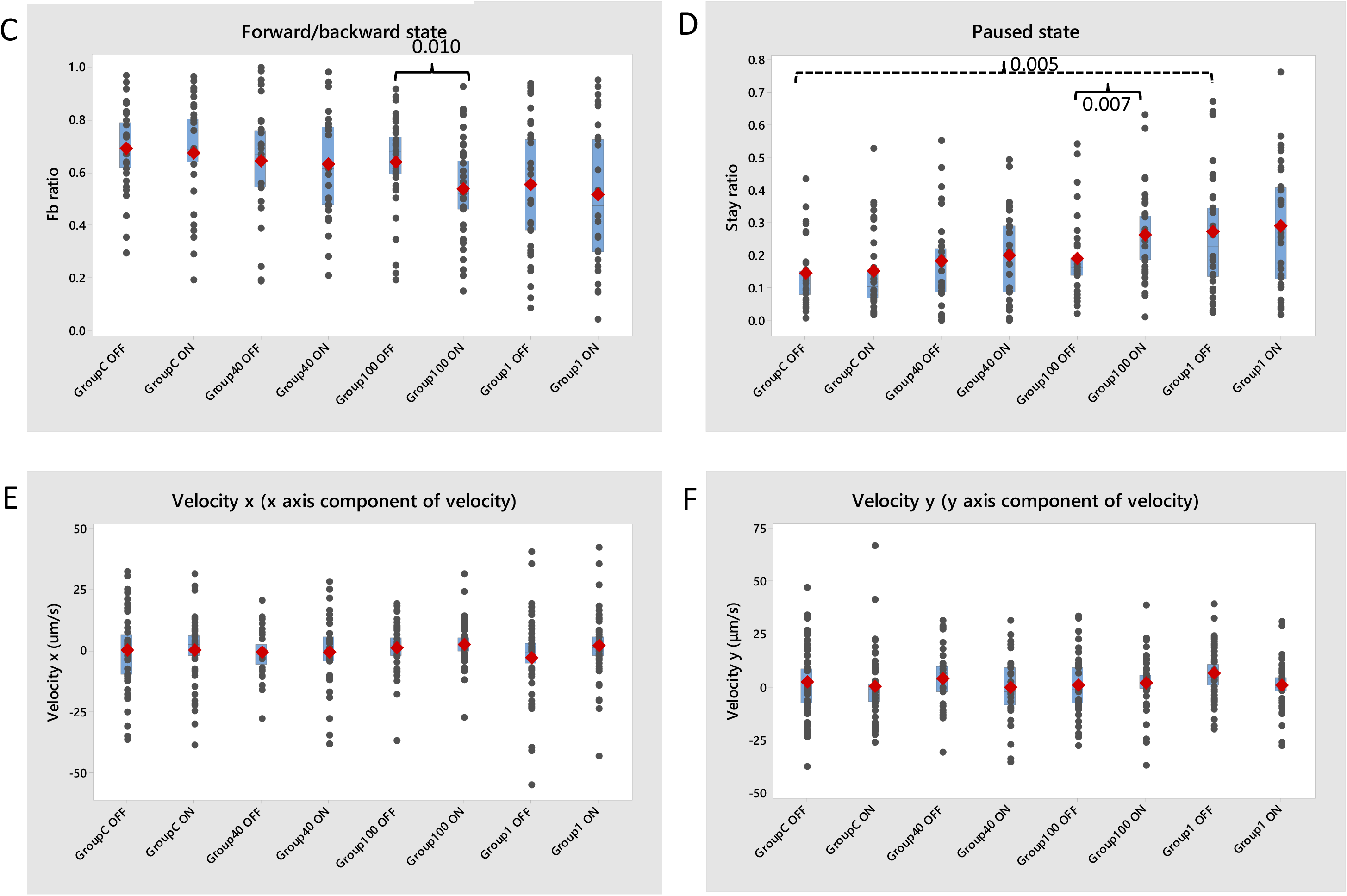
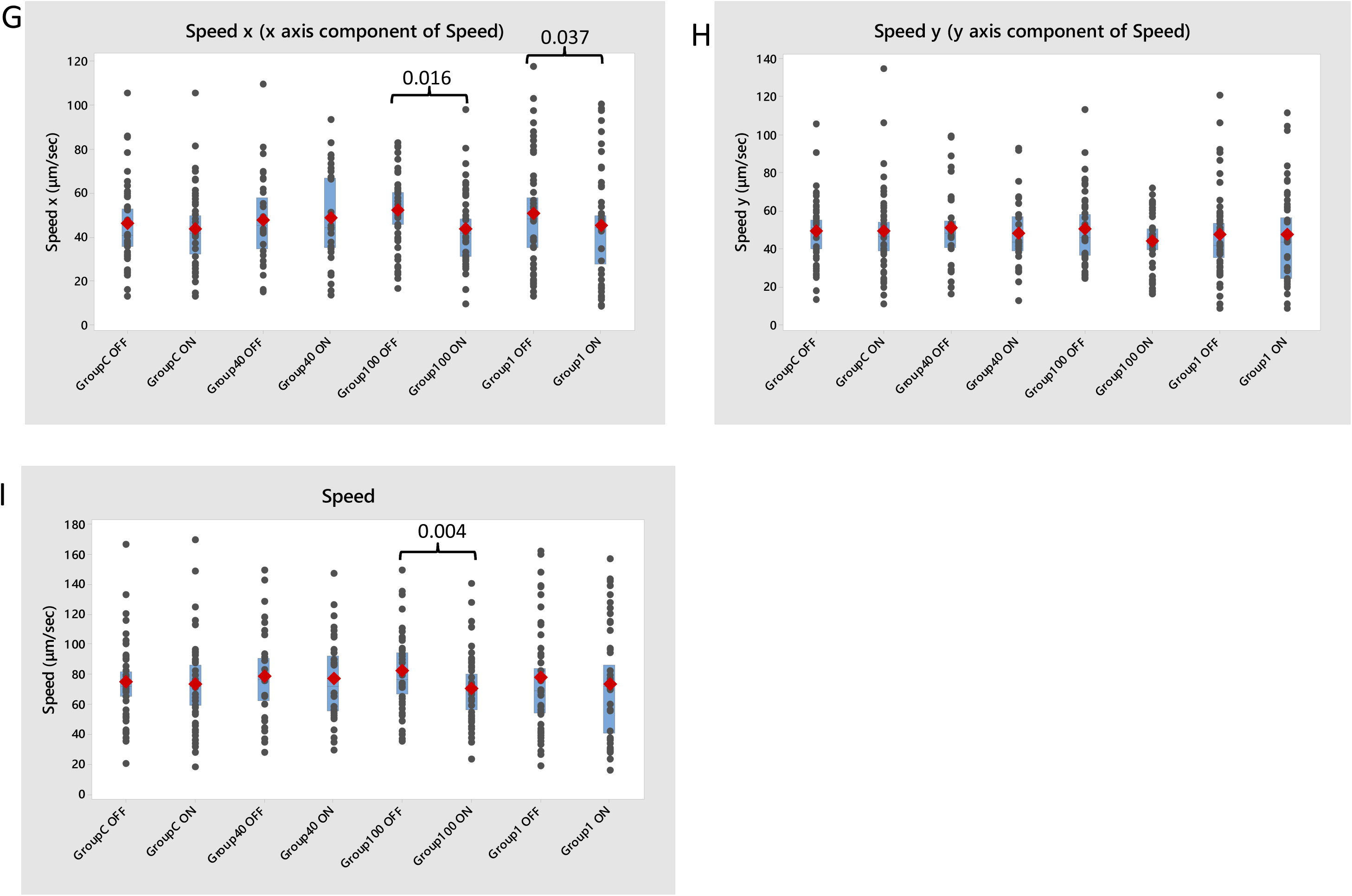
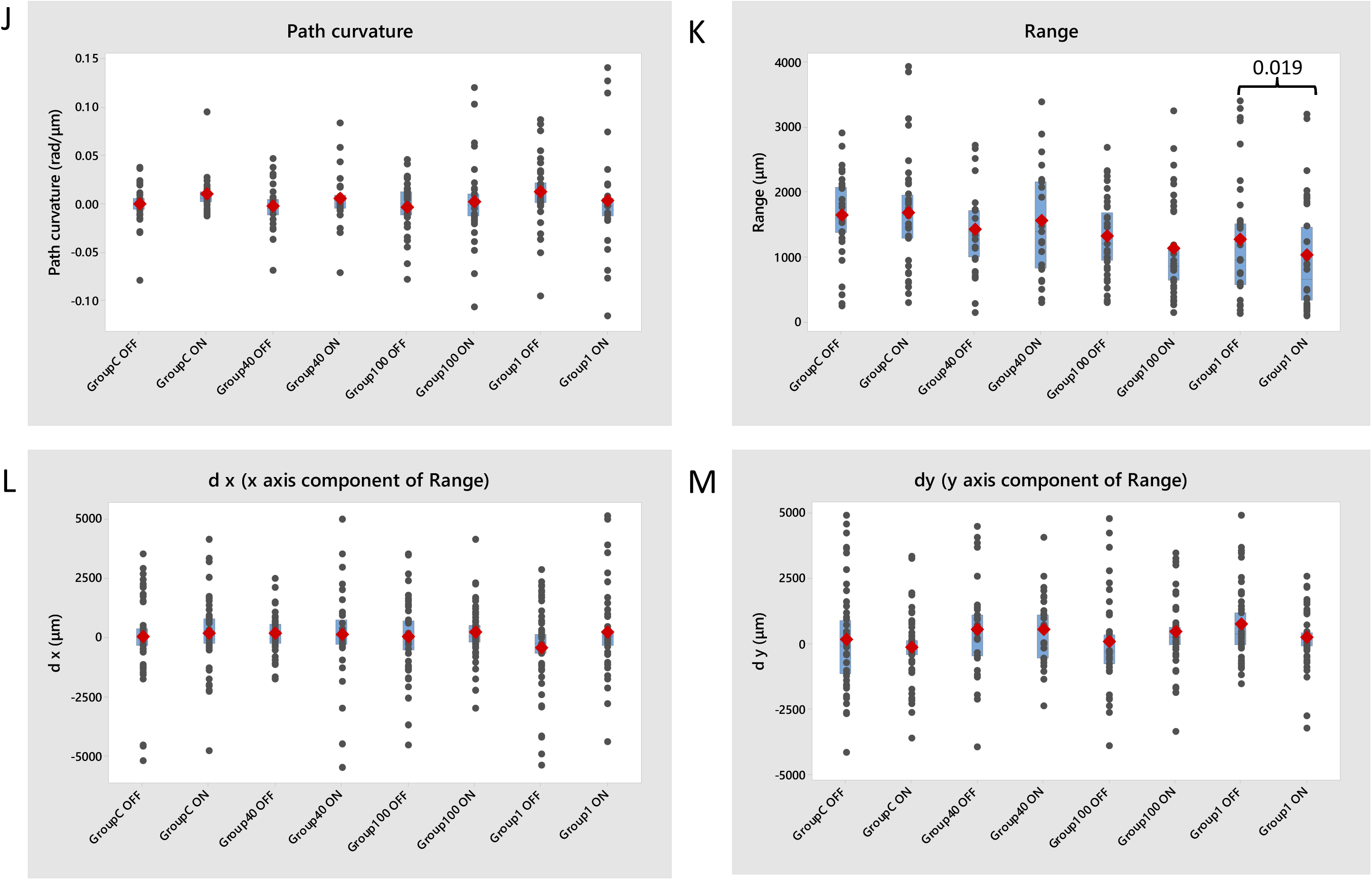
Locomotion features and their metrics, as they have been quantified for all four groups of worms tested. (Group C: control animals, fed on plain food source *E. coli* OP50, n=29; Group 1: fed on *E. coli* OP50 mixed with 1 μm-diameter paramagnetic particles, n=33; Group 100: fed on *E. coli* OP50 mixed with 100 nm-diameter iron core paramagnetic particles, n=38; and Group 40: fed on *E. coli* OP50 mixed with 40 nm-diameter iron core paramagnetic particles, n=42) in the absence (OFF state) or in the presence (ON state) of external magnetic field. A-B: Posture features, C-I: Motion features, J-M: Path features. For each group tested worms were tracked in 3 different experimental days. Grey dots represent individual worms; red diamonds represent the mean; blue boxes indicate the median confidence interval box, with a middle line indicating the median. Dashed lines show comparisons between worms of different groups in the absence of magnetic field (OFF state). Continuous lines show comparisons between the ON and OFF state of worms of the same group. All *p*-values given are calculated by the Wilcoxon Signed Rank test with confidence interval set at 95%, and any difference was considered statistically significant when *p*≤0.05. The *p*-values for all comparisons are given in Tables S1 and S2.

Next, we analyzed features related to the motion state and velocity of the worms. The forward/backward ratio of Group 100 worms decreased when the worms were moving inside the MF (Fig. 6C, WSR test *p*-value = 0.010) and so did their stay ratio (Fig. 6D, WSR test *p*-value = 0.007). Regarding the effect of particles independently of MF, worms of Group 1 had increased stay ratio compared to control animals, even when the MF was turned off (Fig. 6D, Kruskal-Wallis test for all groups in OFF state *p*-value = 0.045, WSR test comparing Group C OFF and Group 1 OFF *p*-value = 0.005). The speed of worms fed with 100 nm particles decreased when MF was on (Fig. 6I, WSR test *p*-value = 0.004), as did speed x for Group 100 (Fig. 6G, WSR test *p*-value = 0.016) and for Group 1 worms (Fig. 6G, WSR test *p*-value = 0.037), whereas speed y component did not change significantly for any group tested (Fig. 6H). Velocity was not affected in any of the groups tested (Fig. 6E and 6F).

We also examined two established path describing features, path curvature and range. The path curvature was not affected by either the presence of particles in the worm body or by MF under the experimental conditions applied (Fig. 6J). However, the range traveled is smaller when worms of Group 1 were moving inside MF compared to the range traveled when worms of the same group were moving without the effect of MF (Fig. 6K, WSR test *p*-value = 0.019). This difference was not reflected in any of the distinct components of range (*d_x_* and *d_y_*, Figs. 6L and 6M, respectively).

The *p*-values for all comparisons are provided in Tables S1 and S2.

## Discussion

### The impact of internalized nanoparticles on *C. elegans* locomotion

Metal nanoparticles of various types have been used to evaluate particle toxicity using *C. elegans* (Gonzalez-Moragas et al., 2017; Gonzalez-Moragas et al., 2015b; Wu et al., 2012; Lim et al., 2012). Particle coating and size, worm developmental stage and duration of exposure have been shown to affect translocation of particles in various tissues of the worm’s body (Pluskota et al., 2009; Gonzalez-Moragas et al., 2017; Wu et al., 2012). Particles used in the present study are larger (Gonzalez-Moragas et al., 2017; Gonzalez-Moragas et al., 2015b; Wu et al., 2012; Lim et al., 2012) and worms have been exposed to them for a shorter period (18-20hrs) than elsewhere (Wu et al., 2012; Yang et al., 2017). These differences may explain why most of the nanoparticles are found along the worms’ pharynx, upper intestine (Fig. 4A, 4B) and lower intestine area (Fig. 4B). The location of 40 nm particles in worms of Group 40 around the pharynx and grinder area (Fig. 4C) is only approximate, as some deformation has been induced on the sample during preparation, and SEM allows for detecting objects that are close to the body surface. Particles aggregates are also found in the intestine (Fig. 5D), intestine lumen (Fig. 5E) and in muscle tissue (Fig. 5F).

Regarding locomotion features, the number of body bends in *C. elegans* L4 larvae has been shown to decrease significantly after exposure for 24 h to 9 nm nanoparticles coated with organic acid (Wu et al., 2012). Worms used in the present work are young adults and not larvae (Pluskota et al., 2009; Yang et al., 2017), which, as developing organisms, could be more vulnerable to toxic effects (Donkin and Williams, 1995). Moreover, particles used in the present work are larger (Gonzalez-Moragas et al., 2017; Gonzalez-Moragas et al., 2015b; Wu et al., 2012; Lim et al., 2012), and made of different metals (Lim et al., 2012; Li et al., 2012), which may result in different ability to overcome the intestine barrier or translocate to other tissues, as well as to different toxicity *per se*. Indeed, in our experiments, exposure of young adults to nanoparticles does not seem to massively affect posture, motion or path features (Fig. 6A–6M).

In our experiments, the only metrics that was affected by nanoparticles alone was the stay ratio for Group 1 (0.25, Fig. 6D), which was higher compared to Group C (0.12, Fig. 6D, comparison indicated by dashed line). This means that Group 1 worms remain paused for longer over the total time recorded, compared to Group C. Aggregates of 1 μm particles in the intestine lumen (Fig. 4A) may resulted in heavier or more cumbersome worms, thus making it more difficult for them to move, although no difference to Group C worms speed and velocity was detected (Fig. 6E–6I). Since there was no MF present, no magnetic effect on the locomotive machinery could take place (see also next section), leaving the locomotion speed unaffected. However, the mass of the particles themselves, or the friction generated between the now heavier worms and the agar surface, could be one possible cause of the observed increased pausing (a single adult *C. elegans* mass is ∼1 μg (Muschiol et al., 2009), and according to Methods, a single worm may have ingested 1 μm nanoparticles up to 10-15% of its body mass). Other effects on *C. elegans* physiology, e.g. impact on muscle function or increased body stiffness, could also be responsible for the increased pausing observed in Group 1 animals. Exploring these issues, however, lies beyond the scope of this paper.

### The impact of internalized nanoparticles, combined with MF, on *C. elegans* locomotion

Magnetotaxis in *C. elegans* was recently demonstrated (Vidal-Gadea, 2015), with the participation of AFD neurons, the first to be identified as magnetosensory. It was suggested that endogenous magnetic material, previously reported in *C. elegans*, may be also involved (Vidal-Gadea, 2015; Cranfield et al., 2004). These findings have sparked an ongoing discussion in the *C. elegans* community (Landler et al., 2018; Vidal-Gadea et al., 2018). In our experiments, the locomotive behavior of Group C worms, which did not contain any particles, was not affected by the externally applied MF (Fig. 6). However, the presence of 100 nm and, in some cases, 1 μm– diameter internalized nanoparticles had an impact on specific locomotion features (Fig. 6), when MF was applied.

Group 100 worms that were moving in MF displayed more body bends (Fig. 6B) and spent more time paused (Fig. 6C, 6D). More body bends possibly indicate a more W-shaped locomotion, which has been described previously in burrowing worms, as opposed to the S-shaped crawling or C-shaped swimming motion (Beron et al., 2015). During burrowing, worms have to put effort to move inside a viscous medium. Hence, one could assume that Group 100 animals crawling in MF integrate more bends to their locomotion, to apply more effort to push their way forward. Indeed, counting the number of body bends has been suggested as a direct measure of the effort a worm is making to move (Hart, 2006). The assumption that moving may be laborious for these animals could be also supported by the fact that they pause more often (Fig. 6C, 6D) and they move more slowly (Fig. 6I). This is observed particularly in the direction of the MF (*x* direction, Fig. 2, Fig. 6G). The magnetic moment for the particles was aligned with the direction of the MF (on the plate surface they both follow the *x* direction, Fig. 3F).

Group 1 worms moving inside MF had also reduced speed in the direction of the MF (*speed_x_*, Fig. 6G), which means that their locomotion was affected especially on the direction parallel to the MF. Moreover, Group 1 worms traveled over a smaller range (Fig. 6K), when the MF was on, which could reflect a modified exploratory behavior (Gray et al., 2005; Cheung et al., 2005). This is more likely to have happened due to changes in the locomotive status related to MF rather than to their incentive to explore, since other environmental factors (e.g., food abundance, temperature) did not change.

It is possible that worms slowed down when they found themselves in a particular orientation inside the MF, or when the internalized particles obtained a particular orientation with regard to the MF. This is supported by the results presented in Figure 2, where it is shown that the properties of MF change significantly in the direction of the MF. Therefore, any effect the MF may had on the particle-fed worm or on the internalized particles themselves, was changing as the worm was moving along the MF.

There was no effect detected in Group 40, in any of the metrics examined. This can be due to the smaller MF or smaller gradient of MF of the particles in this group. However, we have been able to estimate the MF and the gradient of MF only for the 1 µm particles, due to lack of available information on the magnetic properties of the 40 nm and the 100 nm particles. Nonetheless, we can compare the MF and gradient of MF between the particles based on their size (see Supplementary information). We found that smaller particles have larger gradient of MF compared to larger particles in their proximity. The overall impact each particle type has on the worms’ physiology depends on the magnitude and the gradient of MF. Both depend on the material properties of the particles, which determine the magnetic moment. The experimental observations suggested that the stronger effect among the three studied particles occurred in the case of 100 nm particles.

The particles coating (Table 1) was not expected to affect their magnetic behavior. It could, however, impact their interaction with cells. Since the magnetic and physical properties of the particles are the most influential regarding the secondary MF effects, we focused on the particle size for our data analysis. In addition, the thickness of the coating was small (a single monolayer of streptavidin for the 1 μm particles and a 2-3nm thick layer of polymer for the 40 nm particles, according to the manufacturers), which means that the magnetic core could still affect cells and tissues close to the particle. The experimental procedure did not allow us to know the quantity of particles ingested by each individual worm, nor the exact location and the precise interactions of particle aggregates in each individual. Therefore, we cannot extract conclusions based on the particles’ quantity or exact aggregate location inside each worm tested.

Ideally, results shown in Figures 2 and 3 should be combined to assess the synergistic properties of the external component of the MF (generated by the electromagnets) and of the secondary component of the MF (generated by the particles in their vicinity). However, the MF generated by the particles is very localized (in the microscale), as shown in Figure 3. Thus, we discuss below the potential effects of the MF induced by the electromagnets and of the secondary MFs separately, and on their own respective scale. We also discuss the effect of particles aggregates behavior, under the influence of external MF.

### Potential impact of internalized particles, combined with external MF, on *C. elegans* tissues

The forces that were created either by the external MF or the paramagnetic particles themselves were small (Fig. 3E), and they were not strong enough to mechanically push the worms to move along their line of action. Hence, there must be some other mechanism responsible for the detected changes in worms’ locomotion. The magnitude of the external and secondary MFs had the same order of magnitude and they were small (Fig. 2 and 3A, 3C). However, the gradient of the MF fields in the vicinity of the particles was substantially large (Fig. 2B, 2D).

Effects were likely to be more pronounced where the external MF was stronger, since that would result in stronger secondary MFs generated by the particles (until the magnetization of the particles becomes saturated). Hence, the locomotion of worms crawling under stronger MF, namely near the electromagnets (Fig. 2), was more likely to be affected, compared to worms that were moving where the external MF is weaker. Therefore, the spatial distribution of the MF can partly explain the variations observed in the experiments. Moreover, since we had no direct control over the location where particles resided in the worms’ body, the presence or absence of paramagnetic particles where they could affect the mechanosensitive ion channels could be another reason for the variability that we observe in our experiments. In addition, the probability of a worm crawling into areas of higher magnetic flux, would result in it experiencing a stronger effect. The variations in the values of the metrics examined might in fact mirror that probability.

Zablotskii and colleagues (Zablotskii et al., 2016) provide several examples in which the gradient of MF can affect cellular and subcellular mechanisms. The gradient of the secondary MF fields obtained from our simulations (up to 2=10^5^ T/m for the 1 μm particles, Fig. 3B, 3D) was well above the threshold that Zablotskii and colleagues (Zablotskii et al., 2016) suggest may impact cells with mechanosensitive ion channels (10^3^ T/m, see also Table 2 of Zablotskii et al., 2016). The gradient of the secondary internal MF fields was also above the threshold the same authors pose for magnetically induced changes on gene expression, however we consider such a possibility highly unlikely in our case, due to the very short time the external MF was applied (∼ 3.5min). Therefore, the secondary field generated by the paramagnetic particles upon application of external MF could lead to local gradients of MF inside the worm’s body, large enough to interfere with the functionality of excitable cells (e.g., body muscle cells, see also Fig. 5F).

This could have happened by affecting the cells’ ion channels, provided that the particles were very close or even in contact with the cells’ membrane. Experimental data presented here cannot provide insight on exactly which cells might have been the target of the observed MF effect. However, TEM findings (Fig. 5) showed that particles aggregates could be located in body wall muscles (Fig. 5F). Therefore, the possibility that excitable, e.g. muscle cells, were affected by ingested particles in the presence of MF, is considerable. Moreover, impact on the intestine (Fig. 5D, 5E) could affect *C. elegans* physiology and locomotion dynamics, given the multiple roles of this complex tissue (Mcghee, 2007; Nagy et al., 2015). Further experiments are needed to clarify the mechanism behind the observed changes.

The possibility of MF having a direct action on the magnetosensitive neurons described in *C. elegans* by other authors (Vidal-Gadea, 2015) cannot be excluded. However, even if this were true, this action was not reflected in changed locomotion dynamics, which are the object of the present study. Note that in our experiments we did not have direct evidence of sensory or motor neurons being affected by MF, neither was that possibility explored.

Moreover, it is known that when magnetic field is applied to a population of paramagnetic particles, the particles self-organize into arrays, columns, or chains, depending on the nature of the applied magnetic field and the properties of the particle-containing medium (Liu et al., 2005; Liu et al., 1995; Doyle et al., 2002; Mirzakhalili et al., 2017). In our experiments, when the MF is turned ON, it is likely that the internalized particles start moving, as they organize into self-assembled structures. Therefore, it is possible that this motion applies pressure on or stretches the surrounding tissue, resulting in disturbance of the normal locomotion pattern.

### Conclusions

The effect of internalized paramagnetic nanoparticles, in combination with externally applied magnetic field, on the dynamics of *C. elegans*’ locomotion is shown here. Established locomotion metrics, i.e. speed, motion state, bend count, showed differences between untreated worms and worms treated with particles when moving inside magnetic field, while they showed no difference between untreated and particles-treated worms in the absence of magnetic field. Possible explanations on the mechanism that leads to the observed results are provided by work on the effect of magnetic field gradients on cells (Zablotskii et al., 2016; Zablotskii et al., 2014), mediated by magnetic nanoparticles (Hughes et al., 2008), and by work on the self-organizing behavior of paramagnetic particles aggregates inside MF (Liu et al., 2005; Liu et al., 1995; Doyle et al., 2002). The exact mechanism by which the observed effect is achieved in the case of *C. elegans* needs to be further clarified. Our findings on the impact of internalized paramagnetic nanoparticles, in combination with externally applied magnetic field, on animals’ behavior, could pave the way for more detailed studies on the sensitivity of biological systems to these biophysical factors. *C. elegans* nematodes could play a key role in the effort to decipher such phenomena.

## Materials and Methods

### Nanoparticles internalization

i. We investigated the locomotion of four groups of young adult wild type N2 *C. elegans* hermaphrodites, fed on (see also Table 1):
ii. plain bacterial food source *E. coli* OP50, control animals-Group C, n=29;
iii. *E. coli* OP50 mixed with 1 μm-diameter paramagnetic particles (Dynabeads MyOne Streptavidin C1, Invitrogen, Thermo Fisher Scientific, USA), Group 1, n=33;
iv. *E. coli* OP50 mixed with 100 nm-diameter paramagnetic particles (nanomag-CLD-red, Micromod Partikeltechnologie GmbH, Germany), Group 100, n=38 and
v. *E. coli* OP50 mixed with 40 nm-diameter paramagnetic particles (iron oxide nanocrystals, Ocean NanoTech, USA), Group 40, n=42.

In all cases, particles were isolated from the initial suspension by brief centrifugation and were re-suspended in OP50 in a final concentration of 0.5 mg/ml OP50-particle mix. Freshly made 60 mm standard NGM plates were seeded with 100 μl of plain OP50 or OP50-particle mix. Plates were left to dry overnight in room temperature and ∼20 worms were transferred in them the next day. Nematodes were left to feed on the plain or enriched bacterial lawn for 18-20 h, at 20 °C. Then, they were either prepared for microscopy or 12-15 of them were transferred to a fresh, unseeded 35 mm NGM plate for locomotion recording. In the second case, worms were left to acclimatize in the new plate for ∼15 min before recording. There were three reasons for transferring worms to a new, smaller plate. First, we wanted the worms to experience the effect of only internalized nanoparticles under MF and not of the remaining particles on the plate surface. Second, the presence of enriched bacterial lawn on the plate surface interfered with the tracking algorithm and could have affected the worms’ locomotion due to its viscosity. Third, by using 35 mm plates we decreased the distance between the electromagnets and the worms (as shown in Fig. 1), so that the worms experience a stronger external MF.

For each group tested, experiments were run over 3 different experimental days. Therefore, each experimental day we processed 10-14 worms for a specific group. These 10-14 worms were treated simultaneously (on the same plate), and each one of them is considered a biological replicate.

### Fluorescent, Scanning and Transmission Electron Microscopy (SEM and TEM)

#### Fluorescent Microscopy

Worms were transferred to an unseeded NGM plate and were washed with 0.5 ml of 1X PBS. Next, they were transferred to a glass slide, where they were anesthetized on fresh agar pads (Shaham, 2006), using 10 mM NaN_3_ (Sulston, 1988). Samples were imaged using a BX51WI Olympus fluorescent microscope (Olympus, Tokyo, Japan) coupled with an ORCA-flash4.0 camera (Hamamatsu Photonics, Hamamatsu City, Japan).

#### Scanning Electron Microscopy

Samples were prepared as described previously (Hall et al., 1999; Shaham, 2006), with modifications, dissection omitted. Briefly, worms were transferred to an unseeded NGM plate and were washed with 0.5 ml of 1X PBS. Next, they were transferred to a glass cover slip and were anesthetized using 10 mM NaN_3_ (Sulston, 1988). Samples were imaged using FEI Helios 650 nanolab SEM/FIB (FEI, Thermo Fisher Scientific, Waltham, MA).

#### Transmission Electron Microscopy

Samples were prepared based on the literature (Hall et al., 2012; Kovacs, 2015), with modifications. Briefly, tissues were fixed in pre-warmed or RT (room temperature) 3.2% PFA, 0.2% glutaraldehyde in 0.1 M sodium cacodylate buffer for 1h at RT and incubated overnight at 4 °C. Next, they were rinsed 3 × 10 min with 0.1 M sodium cacodylate buffer, and post-fix in 2% OsO4 in 0.1 M sodium cacodylate buffer for 1 h at RT. Another rinse 3 × 10 min with 0.1 M sodium cacodylate buffer followed, and then worms were embedded in resin mold in histogel. Samples were dehydrated for 15 min each in 50%, 70%, 90%, 95%, and finally two changes of 100% ethanol, cleared in two 15 min changes of propylene oxide, and infiltrated in propylene oxide:epon (Embed812), as follows: a.3:1, 1 h, b. 1:1, 1 h, c. 1:3, 1 h, d. Full strength, 2 h or overnight, two changes. Finally, samples were embedded in beam capsules in full epon, and polymerized at 60°C for 24 h. Samples were imaged using a JEM-1400-plus transmission electron microscope (JEOL, Peabody, MA).

### Worm Recording and Tracking

#### Recording

A 35 mm plate containing worms of a specific group was placed between the two electromagnets, as shown in Figure 1, so that the plate surface and therefore the worms were positioned close to the center of the electromagnets. First, a 200 sec movie (1 frame/sec) was recorded in the absence of MF (OFF state) and immediately after, a second 200 sec movie (1 frame/sec) was recorded with the MF on (ON state), using QCapture Pro software, (QImaging, Surrey, Canada) and a Micropublisher3.3 RTV camera (QImaging, Surrey, Canada), mounted on an Olympus SZ61 microscope (Olympus, Tokyo, Japan). We recorded over 200 sec intervals because we were interested in detecting the transient effect of MF on locomotion dynamics. The two electromagnets used were a 4.0′ Dia. Electromagnet, 12 VDC, and a 3.5′ Dia. Electromagnet, 12 VDC, both from APW Company, Rockaway, NJ. Electromagnets were operated at 1.67 A and 3 A respectively, as indicated by the manufacturer, using a 1762 DC power supply (BK Precision, Yorba Linda, CA). By using a non-contact infrared thermometer (Omega Engineering, Norwalk, CT) we verified that the plate surface temperature remained constant throughout the recording period.

#### Tracking

Every movie was imported to MATLAB (MathWorks, Natick, MA) for post-processing. Each worm was tracked individually. To this end, we developed a custom tracking code in MATLAB (Fig. S2). In the first step, all frames were used to construct the movie background, which consisted of all the objects that did not move for long periods of time during the entire recording. Then, each frame was subtracted from the background to extract the foreground, which consisted of all moving objects. Next, the user was prompted with the initial frame of the movie, of which the background had been already subtracted, to select the worm to be tracked by the software. After the user selected the worm, the code created a small examining frame around it and excluded the targeted worm from the rest of the movie frame. Then, the cropped figure was converted to a binary image. After the binary image was enhanced, the shape of the binary object, i.e. the worm, and its global position were stored. Next, the code proceeded to the next movie frame and used the extracted global location of the worm as the center of the small examining frame. The small examining frame must be large enough to capture the motion of the worm in two successive movie frames. Since there was more than one worm freely moving in each experiment, there were occasions in which more than one object were included in the small examining frame. For such occasions, the user was prompted by the code to manually indicate again the worm to be tracked. This way the worm that was initially selected to be tracked was always encapsulated by the examining frame. The code continued the tracking process until the last frame of the movie was processed, and it stored the shape of the worm and its global location for each frame. Once finished, the user run the code again to track another worm.

### Locomotion Analysis

The following features of *C. elegans* morphology and experimental setup properties were used for the quantification of *C. elegans* locomotion parameters.

#### Morphology Features

1. Length: The worm length was defined as the chain-code pixel length of worm skeleton, which was converted into mm.
2. Centroid: The worm density was assumed to be constant throughout its body, so the centroid of mass was the same as geometric centroid. Since the swing of the head or tail (first or last 1/12 chain-code length part of the worm) can significantly influence centroid determination, they were ignored when computing the centroid.

#### Setup

1. Coordinates system: The *x* axis was set along the direction of the MF, between the two electromagnets, and *z* axis was normal to the plate, pointing upwards (Fig.1). Thus, by applying the right-hand rule, the coordinates system was established. Since we did not identify head/tail orientation for the worms, the coordinates system was important for the detection of directionality.
2. Unit Conversion: Any feature regarding length was derived first in pixels. With a known length recorded with the same experimental setup, the conversion between pixels and microns was determined.

Locomotion-related parameters of interest were divided in three categories: posture features, motion features and path features, as described extensively by Yemini and colleagues (Yemini et al., 2013), with minor modifications. A brief description of the examined features follows below.

Posture Features

1. Bends: The total body bends, measured in degrees, derive from the clockwise difference between two tangent supplementary angles (Fig. S4) along the worm skeleton. The mean value (*mean_bends_*) and standard deviation of bends (*std_bends_*) over the worm were also calculated, as intermediate steps.
2. Bend Count: This metric (*bends_num_*) corresponds to the number of bends along a single worm. First, the supplementary angles (see above, Bends) were computed along the worm skeleton. Next, a Gaussian filter over each 1/12 of the chain-code length of the skeleton was applied to the supplementary angles to smooth out any high frequency changes and is then normalized. The filter had a constant proportional to the reciprocal of the standard deviation, α=2.5. By checking the sequence of supplementary angles, the bend count was incremented whenever the angle reaches 0° or changes sign. The check started from the first 1/12 segment to the last 1/12 segment to ignore small bends near the tail and the head.

#### Motion Features

1. Motion State: Worm’s motion state can be divided in two types, the forward/backward state and the paused state. The worm was considered to be in the forward/backward state when its instantaneous speed was greater or equal to 5% of its mean length per second, and it was considered in the paused state when the instantaneous speed was less than 5% of its mean length per second. Therefore, the ratio of the time the worm was in the forward/backward state over the total recording time, namely the *fb_ratio_*, and the ratio of the time the worm was in the paused state over the total recording time, namely the stay ratio *stay_ratio_*, were calculated.
2. Velocity: Velocity is defined as the signed difference between a single worm’s centroids of two sequential frames in the coordinate over the time gap between two frames (1sec). Velocity was further projected on two orthogonal axes *x* and *y* in the plane of the plate (Fig. 1), namely *velocity_x_* and *velocity_y_*. The absolute value of velocity and its components gave speed, *speed_x_* and *speed_y_*, respectively.

#### Path Features

1. Path Curvature: This metric is defined as the angle, in radians, of the worm’s path divided by the distance traveled, in microns. Three successive frames were used to approximate the start, middle and end of the worm’s instantaneous path curvature. The angle was measured by the difference in tangent angles between the second to last frame centroid and the first to the second frame centroid. Then, the path curvature was obtained by dividing the angle by the distance between the first and last centroid.
2. Range: Range is defined as the distance between the worm’s centroid and the centroid of the worm’s path, in each frame. The range was projected onto the orthogonal axes *x* and *y* in the plane of the plate (Fig. 1) to obtain the x-range *d_x_*, and the y-range *d_y_*.

### Magnetic field characterization

COMSOL Multiphysics (COMSOL, Burlington, MA) software was used to characterize the MF that is generated by the two electromagnets in the experimental setup. The data for the magnetic flux density of the electromagnets (available from the manufacturer) was used to calibrate the parameters of the electromagnets in COMSOL Multiphysics. The COMSOL Multiphysics model was used also to estimate the intensity of the external MF, the gradient of the external MF, and the forces that are applied on paramagnetic particles by the external MF. MATLAB was used to calculate the forces applied on the paramagnetic nanoparticles. More details are given in the Supplementary Information section.

### Statistical analysis

Locomotion features were analyzed using non-parametric tests, since the Anderson-Darling normality test *p*-value was > 0.05 for all samples, thus rejecting the normality null hypothesis. For each metric analyzed, the Kruskal-Wallis test was used to detect whether there was any significant difference among the behaviors of all four groups in the absence of MF (OFF state). This comparison was done to determine whether the presence of particles themselves affects locomotion, regardless of external MF. Results were adjusted for ties and any difference was considered statistically significant when *p* ≤ 0.05. To detect differences among worms of the same group, the Wilcoxon Signed Rank (WSR) test was used and differences were considered statistically significant when *p* ≤ 0.05. All analyses were performed in Minitab (Minitab, State College, PA).

To design the experiment, we run a statistical power analysis, using G*Power opensource software (Fig. S5). We prepared the sample size used in the experiments based on this estimation. In order to make sure that we would have enough worms, since some of them might be injured or lost during the process, we slightly increased the sample size number.

## Authors Contributions

BE and EG conceived the idea; EG, EM and BE designed the experiments; EG and YZ run experiments; YZ run tracking algorithm, processed and analyzed recordings; EM created tracking algorithm and run simulations; EG, YZ and EM collected and analyzed data; EG and EM wrote the paper, with input from YZ. All authors reviewed and edited the manuscript and gave final approval for publication.

## Acknowledgments

We thank Nikos Chronis, Kenn Oldham and Jinhong Qu for the use of selected equipment, and Syeda Maisa for help with preliminary worm videos. Scanning electron microscopy was performed at the Electron Microbeam Analysis Laboratory (EMAL), with support from the University of Michigan College of Engineering; we thank John Mansfield, Kai Sun and Haiping Sun for the training. Transmission electron microscopy was performed at the Microscopy & Image Analysis Laboratory (MIL) at the University of Michigan Medical School; we are grateful to Jeff Harrison and Pennelope Blakely for their help and guidance. We thank Surojit Sural for his valuable input on power analysis, and to Hannah Seidel and Elisa Frankel for feedback. We are particularly grateful to Hong Zhan for his generous help with TEM images interpretation.

## Supplementary Information

The features of the magnetic field (MF) presented here are independent of the magnetic particles and depend only on the properties of the electromagnets and the geometry of the setup. However, the forces that are exerted on the particles depend also on the properties of the particle. We have used several different magnetic particles in our experiments. The numerical simulations and analysis are performed only on the largest magnetic particles since such analysis provide us with the largest forces that can be created because of the external MF. Moreover, the characteristics of Dynabeads are more accessible (Fonnum et al., 2005) compared to the rest of the particles used in our experiments.

The forces on a magnetic particle in an external MF can be characterized as (Shevkoplyas et al., 2007):

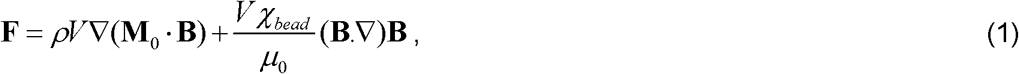

where *ρ*, *V*, *M*_0_, *χ_bead_* are the density, volume, initial magnetization, and initial magnetic susceptibility of the particles, respectively. **F** is the force acting on the particle, **B** is the external MF, and *μ*_0_ is the permeability of vacuum.

Equation (1) is used with the properties of the magnetic particles (Fonnum et al., 2005) and the simulation results from COMSOL Multiphysics for the external MF to calculate the forces acting on the magnetic particles. Figure 2 shows that the magnitude of the force at all locations on the plate, which, as expected, can be observed to be larger near the electromagnets where the MF and its gradient are larger.

The magnetization of the particles creates local modifications to the MF compared to the externally applied MF. From classical physics, the MF around a magnetic dipole can be expressed as:

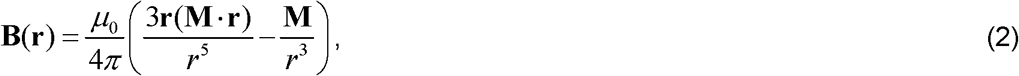

where **r** is position vector relative to the particle at which the MF is calculated and **M** is the magnetic moment of the particle that can be calculated as (Shevkoplyas et al., 2007):

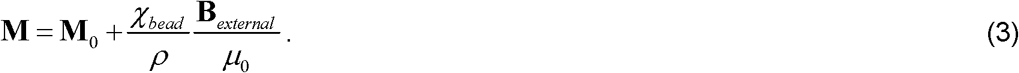

Equation (3) is used with the properties of the largest particles in our experiments (Fonnum et al., 2005) along with the external MF (**B***_external_*) created by the electromagnets obtained from COMSOL Multiphysics to determine the magnetic moments of the particles in the presence of the external MF. Next Eq. (2) is used to calculate the MF around the particles when the magnetic moment is known. Figure 3 shows the magnitude of the MF flux for vertical and horizontal configuration of three magnetic particles using Eqs. (2) and (3) in MATLAB. The MF is strong close to the particles for both configurations, and decays rapidly with distance from the particles.

The MF shown in Figure 3 can also be used to calculate the gradient of the MF around the particles. Figure 3 show the gradient of the magnetic of for both horizontal and vertical configuration. Again, the gradient of the MF is strongest near the particles and decays rapidly with distance from the particles.

Finally, the forces between two magnetic particles can be found as (3):

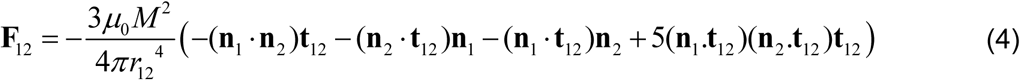

Where **F**_12_ is the force between the two particles, *r*_12_ is the distance between the two particles, **t**_12_ is the unit vector that connects the two particles, **n**_1_ and **n**_2_ are the direction of the magnetic moment for particle 1 and 2, and *M* is the magnitude of the magnetic moment of the two the same magnetic moment for both particles based on Eq. (3). Figure 3E shows how the force between two particles changes for both vertical and horizontal direction. The force between the particles in the horizontal direction is repulsive while the force between the particles in the vertical direction is attractive. The attractive forces between the particles cause them to form chain-like structures when they are not interrupted by the medium in which the particles are located (Mirzakhalili et al., 2017; Nakata et al., 2008).

### Comparison of the MF/gradient MF between particles

We compare the MF and gradient of MF between the particles based on their size. The iron mass of an ideal spherical particle scales with *d*^3^, where *d* is the diameter of the particle. Hence, for constant density, smaller particles have smaller magnetic mass by a factor of *d*^−3^. However, according to Eq. 2 in the Supplementary Information, the MF around a particle scales with *r*^−3^, where *r* is the distance from the center of the particle. Therefore, the MF on the surface of a particle changes with its size. Smaller particles have less magnetic material, so their MF in the proximity of the particle is smaller. In addition, the gradient of MF scales with *r*^−4^. Hence, since the mass of the magnetic core scales with *d*^−3^, the gradient of MF on the surface of the particle scales with *d*. Therefore, smaller particles will have larger gradient of MF compared to larger particles in their proximity.

---

**Fig. S1:**
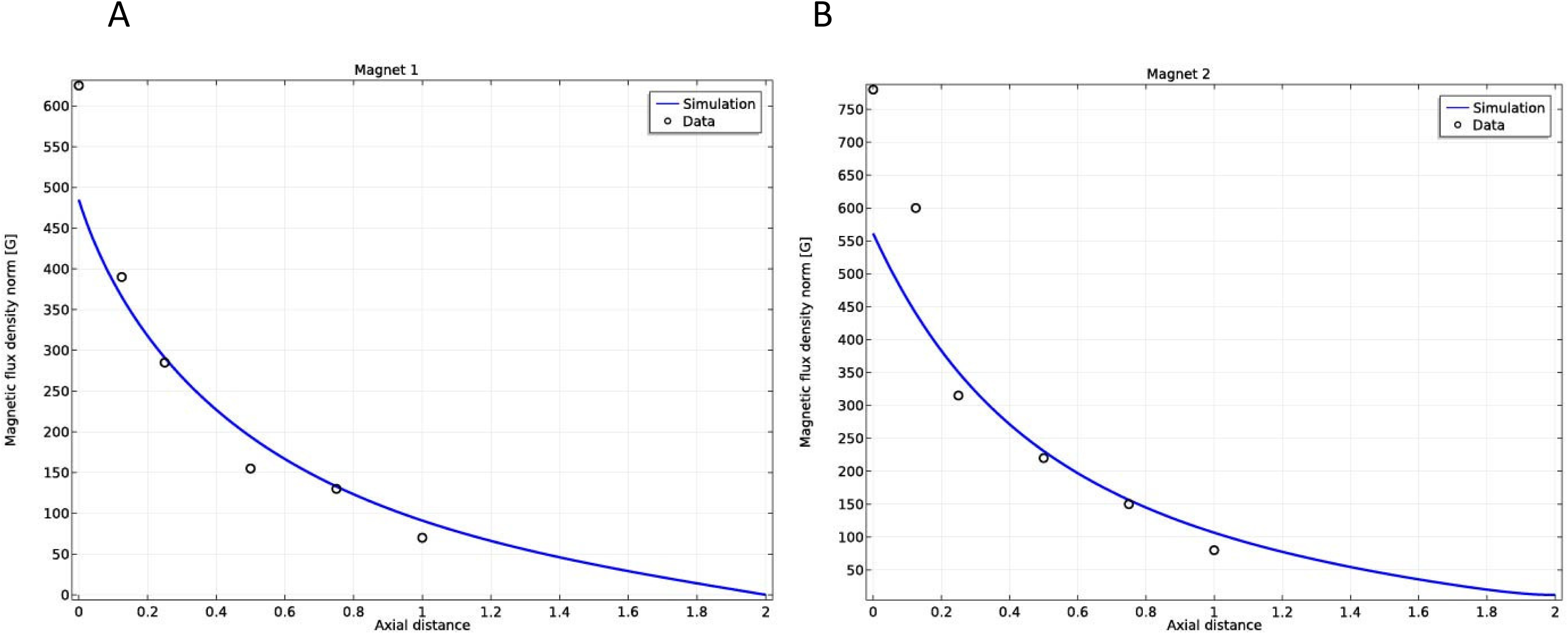
Calibration of the parameters in COMSOL Multiphysics simulations to match the available data for magnet 1 (A) and magnet 2 (B) that are used in the experiments.

**Fig. S2:**
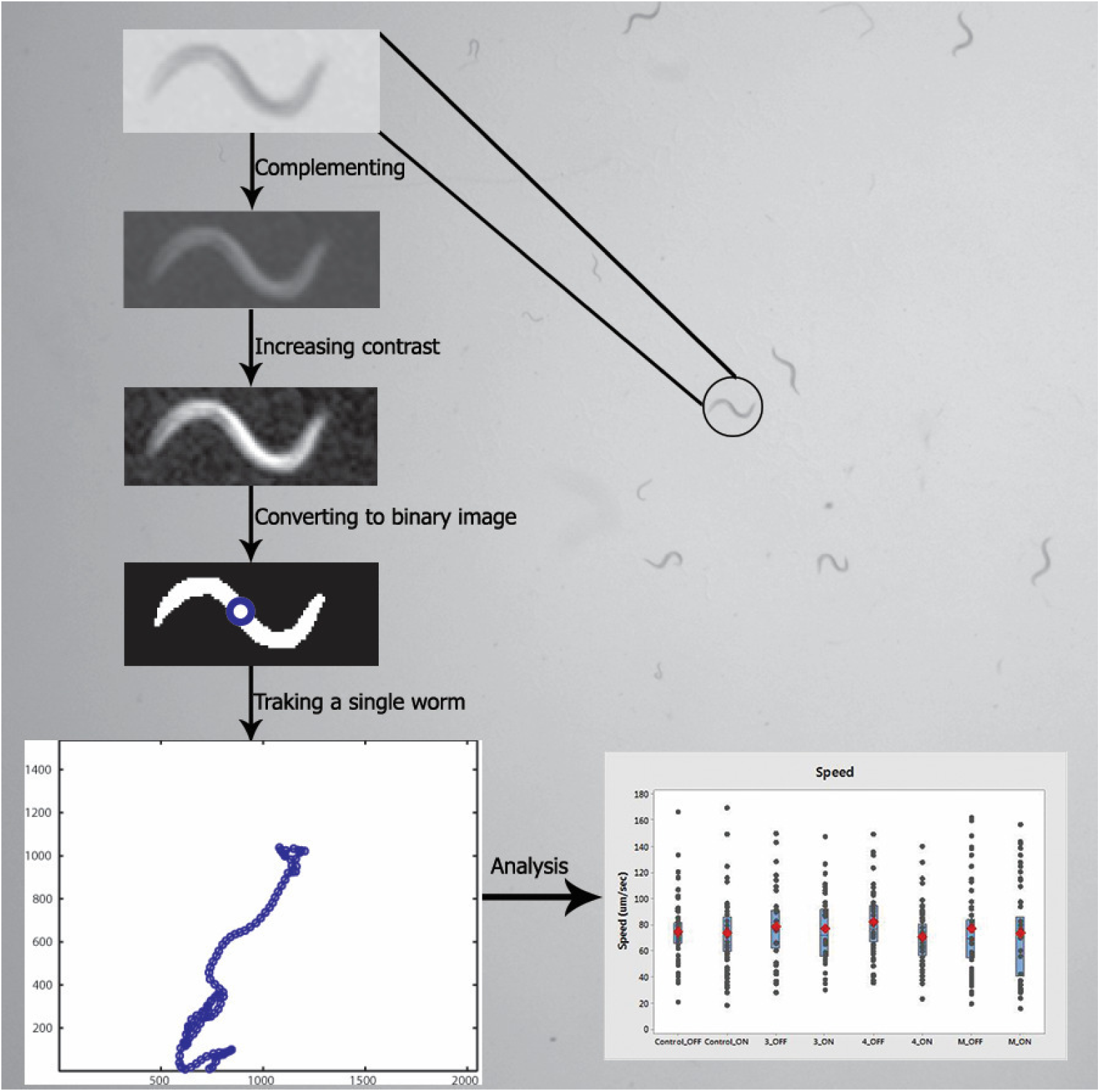
An overview of the steps that are taken to analyze the locomotion of the worms. Top to bottom: A worm is selected in the first frame to be tracked. Next, several image enhancements are performed on the subframe that is created around the selected worm. Next, the grayscale image is converted to a binary image and postprocessing, e.g. finding the centroid (blue circle) of the worm, is performed. All these steps are conducted for the whole movie that is recorded. Finally, the data collected from all experiments are compared.

**Fig. S3:**
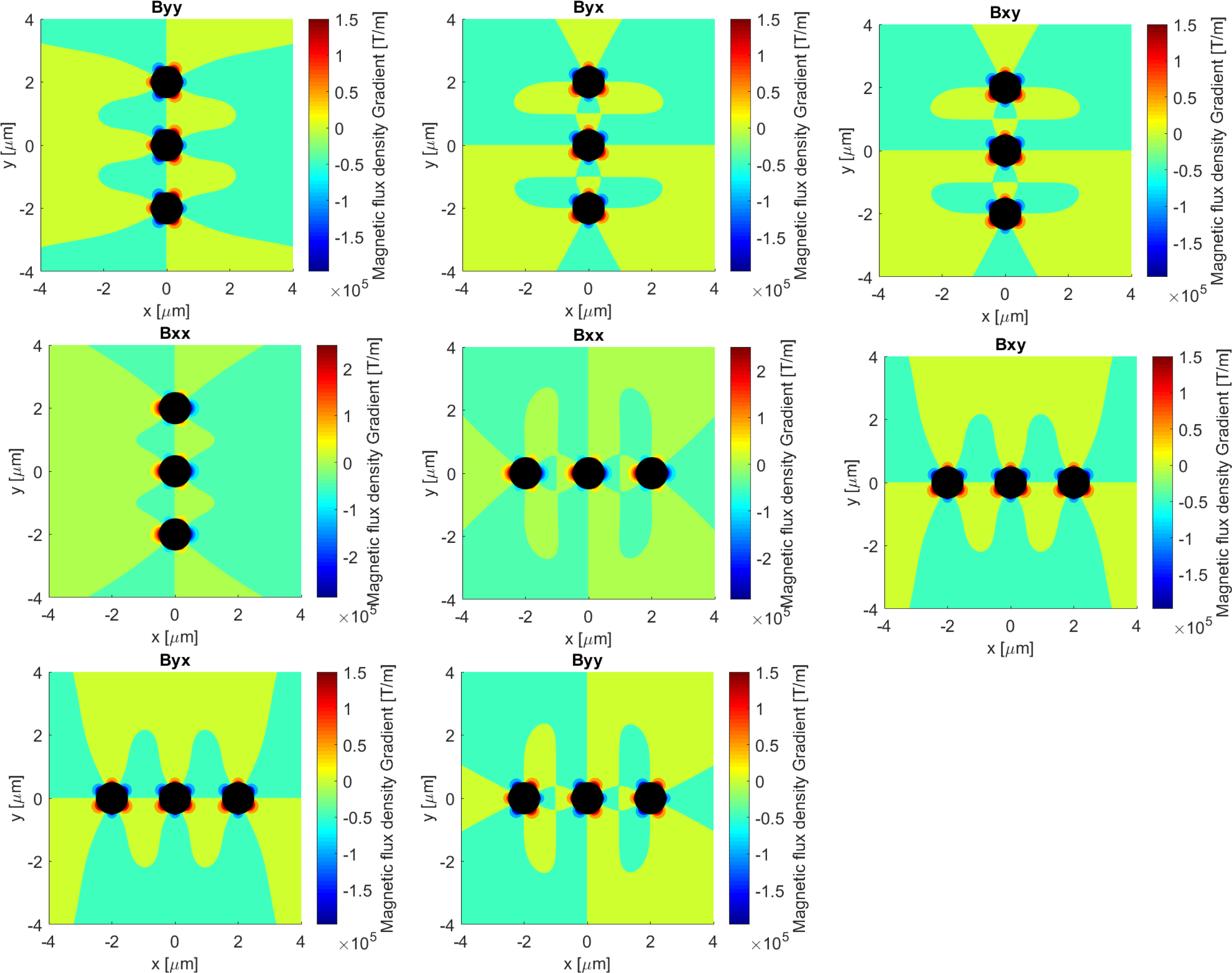
All the components of the gradient of the magnetic field for the particles in the vertical (along the y axis) and horizontal (along the x axis) configuration.

**Fig. S4:**
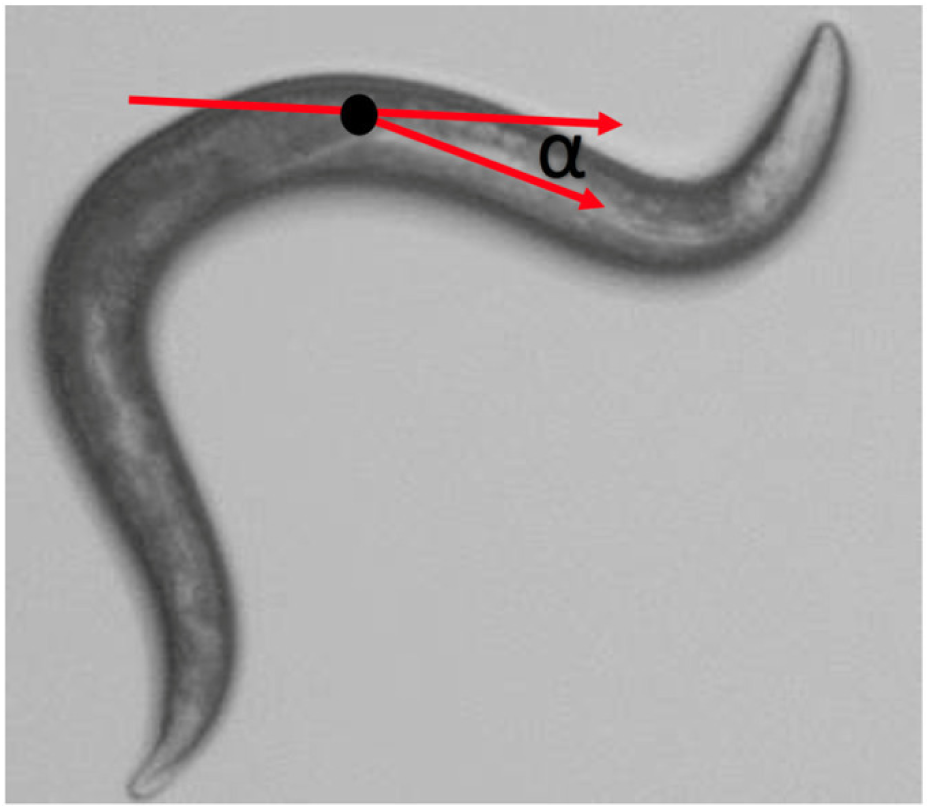
The supplementary angle (α) is the difference in tangent angles at each skeleton point.

**Table S1:** The magnetic field effect on the locomotion of worms with internalized nanoparticles. *p*-values of Wilcoxon Signed Rank test for all comparisons between OFF and ON state of each group studied. With bold are highlighted the statistically significant differences, with *p*-value ≤0.05.

| Metric | Group C OFF/ON | Group 1 OFF/ON | Group 100 OFF/ON | Group 40 OFF/ON |
| --- | --- | --- | --- | --- |
| Total bends | 0.645 | 0.645 | 0.644 | 0.540 |
| Bend count | 0.052 | 0.093 | **0.031** | 0.764 |
| Forward/backward ratio | 0.681 | 0.193 | **0.010** | 0.747 |
| Stay ratio | 0.967 | 0.336 | **0.007** | 0.706 |
| Velocity x | 0.832 | 0.375 | 0.477 | 0.979 |
| Velocity y | 0.599 | 0.066 | 0.514 | 0.375 |
| Speed x | 0.431 | **0.037** | **0.016** | 0.936 |
| Speed y | 0.235 | 0.066 | 0.093 | 0.435 |
| Speed | 0.438 | 0.190 | **0.004** | 0.554 |
| Path curvature | 0.120 | 0.059 | 0.543 | 0.225 |
| Range | 0.773 | **0.019** | 0.104 | 0.706 |
| D x | 0.960 | 0.280 | 0.499 | 0.914 |
| D y | 0.626 | 0.250 | 0.217 | 0.920 |

**Table S2:** The particle effect on the locomotion of worms with internalized nanoparticles. In the second column are given the *p*-values of Kruskal-Wallis test used to compare all groups of worms studied during their OFF state. In the last three columns are given the *p*-values of the Wilcoxon Signed Rank test used to compare Group C with each of the other three particle-containing groups. The Wilcoxon Signed Rank test was run only when Kruskal-Wallis test gave a *p*-value ≤0.05. With bold are highlighted the statistically significant differences, with *p*-value ≤0.05.

| Metric | Kruskal-Wallis | Group C vs Group 1 | Group C vs Group 100 | Group C vs Group 40 |
| --- | --- | --- | --- | --- |
| Total bends | 0.649 |  |  |  |
| Bend count | 0.237 |  |  |  |
| Forward/backward ratio | 0.234 |  |  |  |
| Stay ratio | **0.045** | **0.005** | 0.226 | 0.064 |
| Velocity x | 0.729 |  |  |  |
| Velocity y | 0.321 |  |  |  |
| Speed x | 0.498 |  |  |  |
| Speed y | 0.804 |  |  |  |
| Speed | 0.661 |  |  |  |
| Path curvature | 0.096 |  |  |  |
| Range | 0.115 |  |  |  |
| D x | 0.649 |  |  |  |
| D y | 0.188 |  |  |  |

**Fig. S5:**
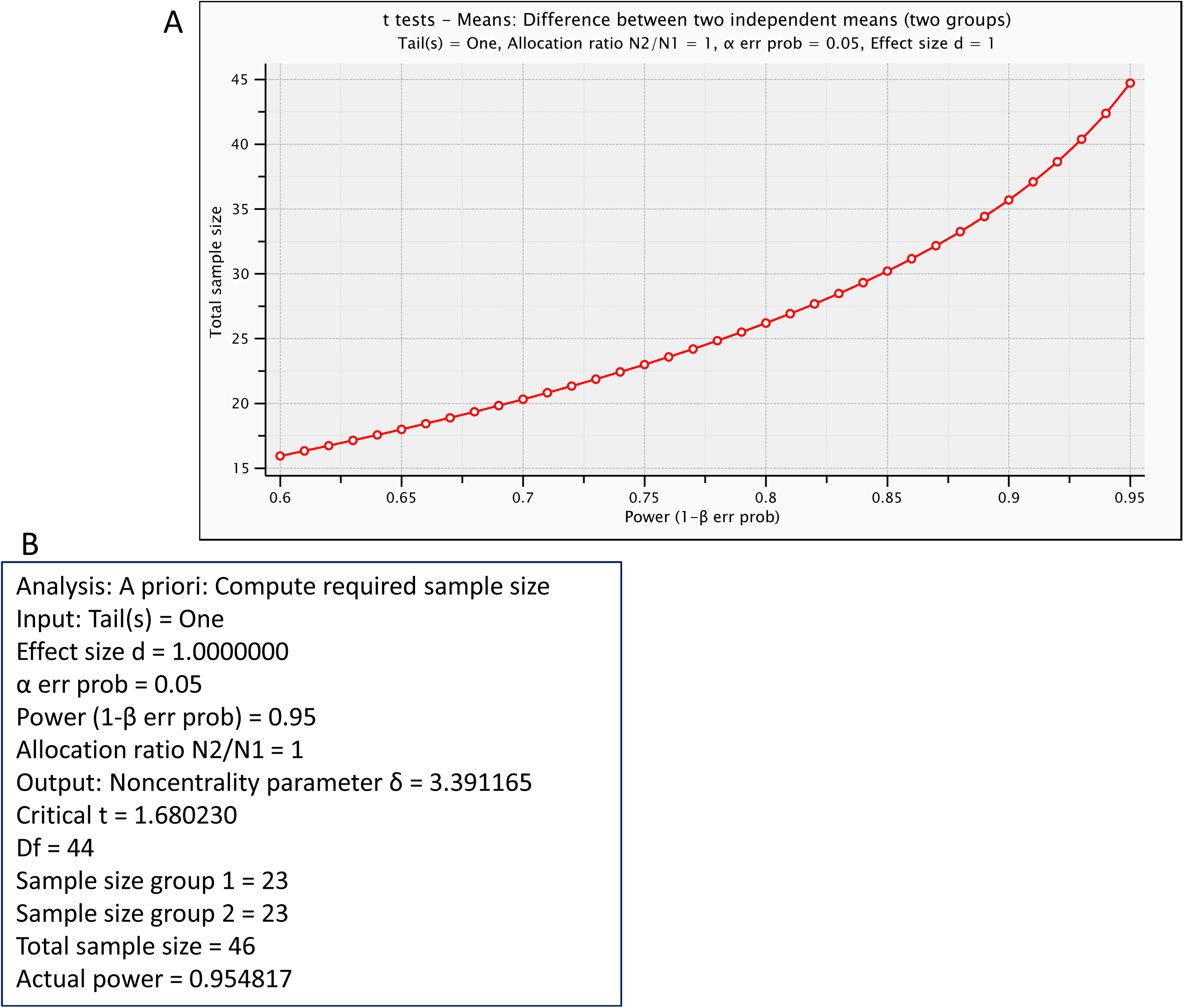
Power analysis for locomotion dynamics experiments. G*Power open source software was used as described in Methods. A. Plot that shows the probability of detecting a real effect with regard to sample size; B. Table showing the protocol followed for the power analysis. The mean of group1 was set to 0.5 and the mean of group2 was set to 0.4, with SD within each group σ=0.1.

## References

Bae, J.-E., Bang, S., Min, S., Lee, S.-H., Kwon, S.-H., Lee, Y., Lee, Y.-H., Chung, J. & Chae, K.-S. 2016. Positive geotactic behaviors induced by geomagnetic field in Drosophila. Molecular Brain, 9, 55.

Bansal, A., Zhu, L. J., Yen, K. & Tissenbaum, H. A. 2015. Uncoupling lifespan and healthspan in Caenorhabditis elegans longevity mutants. Proceedings of the National Academy of Sciences, 112, E277–E286.

Belova, N. A. & Acosta-Avalos, D. 2015. The Effect of Extremely Low Frequency Alternating Magnetic Field on the Behavior of Animals in the Presence of the Geomagnetic Field. Journal of Biophysics, 2015, 8.

Beron, C., Vidal-Gadea, A. G., Cohn, J., Parikh, A., Huong, G. & Pierce-Shimomura, J. T. 2015. The burrowing behavior of the nematode Caenorhabditis elegans: A new assay for the study of neuromuscular disorders. Genes, brain, and behavior, 14, 357–368.

Cheung, B. H., Cohen, M., Rogers, C., Albayram, O. & De Bono, M. 2005. Experience-dependent modulation of C. elegans behavior by ambient oxygen. Current Biology, 15, 905–917.

Cranfield, C. G., Dawe, A., Karloukovski, V., Dunin-Borkowski, R. E., De Pomerai, D. & Dobson, J. 2004. Biogenic magnetite in the nematode caenorhabditis elegans. Proceedings in Biological Sciences, 271 Suppl 6, S436–S439.

De Bono, M. & Bargmann, C. I. 1998. Natural Variation in a Neuropeptide Y Receptor Homolog Modifies Social Behavior and Food Response in C. elegans. Cell, 94, 679–689.

Donkin, S. G. & Williams, P. L. 1995. Influence of developmental stage, salts and food presence on various end points using Caenorhabditis Elegans for aquatic toxicity testing. Environmental Toxicology and Chemistry, 14, 2139–2147.

Doyle, P. S., Bibette, J., Bancaud, A. & Viovy, J.-L. 2002. Self-Assembled Magnetic Matrices for DNA Separation Chips. Science, 295, 2237–2237.

Fedele, G., Green, E. W., Rosato, E. & Kyriacou, C. P. 2014. An electromagnetic field disrupts negative geotaxis in Drosophila via a CRY-dependent pathway. Nature Communications, 5, 4391.

Ghodbane, S., Lahbib, A., Sakly, M. & Abdelmelek, H. 2013. Bioeffects of static magnetic fields: oxidative stress, genotoxic effects, and cancer studies. Biomedical Research International 2013, 602987.

Giachello, C. N. G., Scrutton, N. S., Jones, A. R. & Baines, R. A. 2016. Magnetic Fields Modulate Blue-Light-Dependent Regulation of Neuronal Firing by Cryptochrome. The Journal of Neuroscience, 36, 10742–10749.

Gonzalez-Moragas, L., Roig, A. & Laromaine, A. 2015a. C. elegans as a tool for in vivo nanoparticle assessment. Advances in Colloid and Interface Science, 219, 10–26.

Gonzalez-Moragas, L., Yu, S.-M., Benseny-Cases, N., Stürzenbaum, S., Roig, A. & Laromaine, A. 2017. Toxicogenomics of iron oxide nanoparticles in the nematode C. elegans. Nanotoxicology, 11, 647–657.

Gonzalez-Moragas, L., Yu, S.-M., Carenza, E., Laromaine, A. & Roig, A. 2015b. Protective Effects of Bovine Serum Albumin on Superparamagnetic Iron Oxide Nanoparticles Evaluated in the Nematode Caenorhabditis elegans. ACS Biomaterials Science & Engineering, 1, 1129–1138.

Gourgou, E., Chronis, N. 2016. Chemically induced oxidative stress affects ASH neuronal function and behavior in C. elegans. Scientific Reports, 6, 38147.

Gray, J. M., Hill, J. J. & Bargmann, C. I. 2005. A circuit for navigation in Caenorhabditis elegans. Proceedings of the National Academy of Sciences of the United States of America, 102, 3184–3191.

Hahm, J. H., Kim, S., Diloreto, R. & Shi, C. 2015. C. elegans maximum velocity correlates with healthspan and is maintained in worms with an insulin receptor mutation. Nature Communications, 6, 8919.

Hall, D. H., Hartwieg, E. & Nguyen, K. C. 2012. Modern electron microscopy methods for C. elegans. Methods in Cell Biology, 107, 93–149.

Hall, D. H., Winfrey, V. P., Blaeuer, G., Hoffman, L. H., Furuta, T., Rose, K. L., Hobert, O. & Greenstein, D. 1999. Ultrastructural Features of the Adult Hermaphrodite Gonad of Caenorhabditis elegans: Relations between the Germ Line and Soma. Developmental Biology, 212, 101–123.

Hart, A. C. 2006. Behavior. In: Ambros (ed.) Wormbook. The C. elegans Research Community.

Hedgecock, E. M. & Russell, R. L. 1975. Normal and mutant thermotaxis in the nematode Caenorhabditis elegans. Proceedings of the National Academy of Sciences of the United States of America 72, 4061–4065.

Hong, F. T. 1995. Magnetic field effects on biomolecules, cells, and living organisms. Biosystems, 36, 187–229.

Hsu, A.-L., Feng, Z., Hsieh, M.-Y. & Xu, X. Z. S. 2009. Identification by machine vision of the rate of motor activity decline as a lifespan predictor in C. elegans. Neurobiology of aging, 30, 1498–1503.

Huang, H., Delikanli, S., Zeng, H., Ferkey, D. M. & Pralle, A. 2010. Remote control of ion channels and neurons through magnetic-field heating of nanoparticles. Nature Nanotechnology 5, 602–606.

Hughes, S., Mcbain, S., Dobson, J. & El Haj, A. J. 2008. Selective activation of mechanosensitive ion channels using magnetic particles. Journal of Royal Society Interface, 5, 855–863.

Kale, P. G. & Baum, J. W. 1980. Genetic effects of strong magnetic fields in drosophila melanogaster: II. Lack of interaction between homogeneous fields and fission neutron-plus-gamma radiation. Environmental Mutagenesis, 2, 179–186.

Khare, P., Sonane, M., Pandey, R., Ali, S., Gupta, K. C. & Satish, A. 2011. Adverse Effects of TiO2 and ZnO Nanoparticles in Soil Nematode, Caenorhabditis elegans. Journal of Biomedical Nanotechnology, 7, 116–117.

Kim, J. H., Lee, S. H., Cha, Y. J., Hong, S. J., Chung, S. K., Park, T. H. & Choi, S. S. 2017. C. elegans-on-a-chip for in situ and in vivo Ag nanoparticles’ uptake and toxicity assay. Scientific Reports, 7, 40225.

Kovacs, A. L. 2015. The application of traditional transmission electron microscopy for autophagy research in Caenorhabditis elegans. Biophys Rep, 1, 99–105.

Kumari, K., Capstick, M., Cassara, A. M., Herrala, M., Koivisto, H., Naarala, J., Tanila, H., Viluksela, M. & Juutilainen, J. 2017. Effects of intermediate frequency magnetic fields on male fertility indicators in mice. Environmental Research, 157, 64–70.

Landler, L., Nimpf, S., Hochstoeger, T., Nordmann, G. C., Papadaki-Anastasopoulou, A. & Keays, D. A. 2018. Comment on “Magnetosensitive neurons mediate geomagnetic orientation in Caenorhabditis elegans”. eLife, 7, e30187.

Lee, C.-H., Hung, Y.-C. & Huang, G. S. 2010. Static magnetic field accelerates aging and development in nematode. Communicative & Integrative Biology, 3, 528–529.

Lewczuk, B., Redlarski, G., Arkadiusz, Zikowska, N., Przybylska-Gornowicz, B., Krawczuk, M. 2014. Influence of Electric, Magnetic, and Electromagnetic Fields on the Circadian System: Current Stage of Knowledge. BioMed Research International, 2014, 13.

Li, G., Gong, J., Lei, H., Liu, J. & Xu, X. Z. S. 2016. Promotion of behavior and neuronal function by reactive oxygen species in C. elegans. Nature Communications, 7, 13234.

Li, Y., Yu, S., Wu, Q., Tang, M., Pu, Y. & Wang, D. 2012. Chronic Al2O3-nanoparticle exposure causes neurotoxic effects on locomotion behaviors by inducing severe ROS production and disruption of ROS defense mechanisms in nematode Caenorhabditis elegans. J Hazard Mater, 219-220, 221–230.

Liedtke, W., Tobin, D. M., Bargmann, C. I. & Friedman, J. M. 2003. Mammalian TRPV4 (VR-OAC) directs behavioral responses to osmotic and mechanical stimuli in Caenorhabditis elegans. Proceedings of the National Academy of Sciences, 100, 14531–14536.

Lim, D., Roh, J. Y., Eom, H. J., Choi, J. Y., Hyun, J. & Choi, J. 2012. Oxidative stress-related PMK-1 P38 MAPK activation as a mechanism for toxicity of silver nanoparticles to reproduction in the nematode Caenorhabditis elegans. Environ Toxicol Chem, 31, 585–592.

Liu, D., Maxey, M. R. & Karniadakis, G. E. 2005. Simulations of dynamic self-assembly of paramagnetic microspheres in confined microgeometries. Journal of Micromechanics and Microengineering, 15, 2298.

Liu, J., Lawrence, E. M., Wu, A., Ivey, M. L., Flores, G. A., Javier, K., Bibette, J. & Richard, J. 1995. Field-Induced Structures in Ferrofluid Emulsions. Phys Rev Lett, 74, 2828–2831.

Liu, J., Zhang, B., Lei, H., Feng, Z., Liu, J., Hsu, A. L., Xu, X. Z. 2013. Functional aging in the nervous system contributes to age-dependent motor activity decline in C. elegans. Cell Metab, 18, 392–402.

Long, X., Ye, J., Zhao, D. & Zhang, S.-J. 2015. Magnetogenetics: remote non-invasive magnetic activation of neuronal activity with a magnetoreceptor. Science Bulletin, 60, 2107–2119.

Ma, H., Bertsch, P. M., Glenn, T. C., Kabengi, N. J. & Williams, P. L. 2009. Toxicity of manufactured zinc oxide nanoparticles in the nematode Caenorhabditis elegans. Environ Toxicol Chem, 28, 1324–1330.

Malkemper, E. P., Eder, S. H. K., Begall, S., Phillips, J. B., Winklhofer, M., Hart, V. & Burda, H. 2015. Magnetoreception in the wood mouse (Apodemus sylvaticus): influence of weak frequency-modulated radio frequency fields. Scientific Reports, 5, 9917.

Mcghee, J. D. 2007. The C. elegans intestine. In: Seydoux G. (ed.) WormBook. The C. elegans Research Community.

Meyer, J. N., Lord, C. A., Yang, X. Y., Turner, E. A., Badireddy, A. R., Marinakos, S. M., Chilkoti, A., Wiesner, M. R. & Auffan, M. 2010. Intracellular uptake and associated toxicity of silver nanoparticles in Caenorhabditis elegans. Aquat Toxicol, 100, 140–150.

Mirzakhalili, E., Nam, W. & Epureanu, B. I. 2017. Reduced-order models for the dynamics of superparamagnetic nanoparticles interacting with cargoes transported by kinesins. Nonlinear Dynamics, 90, 425–442.

Miyakoshi, J. 2005. Effects of static magnetic fields at the cellular level. Prog Biophys Mol Biol, 87, 213–223.

Muschiol, D., Schroeder, F. & Traunspurger, W. 2009. Life cycle and population growth rate of Caenorhabditis elegans studied by a new method. BMC Ecology, 9, 14–14.

Nagy, S., Huang, Y.-C., Alkema, M. J. & Biron, D. 2015. Caenorhabditis elegans exhibit a coupling between the defecation motor program and directed locomotion. Scientific Reports, 5, 17174.

Naito, M., Hirai, S., Mihara, M., Terayama, H., Hatayama, N., Hayashi, S., Matsushita, M. & Itoh, M. 2012. Effect of a Magnetic Field on Drosophila under Supercooled Conditions. PLoS ONE, 7, e51902.

Nakata, K., Hu, Y., Uzun, O., Bakr, O. & Stellacci, F. 2008. Chains of Superparamagnetic Nanoparticles. Advanced Materials, 20, 4294–4299.

Njus, Z., Feldmann, D., Brien, R., Kong, T., Kalwa, U., Pandey, S. 2015. Characterizing the Effect of Static Magnetic Fields on *C. elegans* Using Microfluidics. Advances in Bioscience and Biotechnology, 06, 583–591.

Öcal, I., Kalkan, T., Ganay, O. 2008. Effects of alternating magnetic field on the metabolism of the healthy and diabetic organisms. Brazilian Archives of Biology and Technology, 51, 523–530.

Osipova, E. A., Pavlova, V. V., Nepomnyashchikh, V. A. & Krylov, V. V. 2016. Influence of magnetic field on zebrafish activity and orientation in a plus maze. Behav Processes, 122, 80–86.

Parida, L., Neogi, S. & Padmanabhan, V. 2014. Effect of Temperature Pre-Exposure on the Locomotion and Chemotaxis of C. elegans. PLOS ONE, 9, e111342.

Peliti, M., Chuang, J. S. & Shaham, S. 2013. Directional Locomotion of C. elegans in the Absence of External Stimuli. PLOS ONE, 8, e78535.

Pierce-Shimomura, J. T., Chen, B. L., Mun, J. J., Ho, R., Sarkis, R. & Mcintire, S. L. 2008. Genetic analysis of crawling and swimming locomotory patterns in C. elegans. Proceedings of the National Academy of Sciences of the United States of America, 105, 20982–20987.

Pluskota, A., Horzowski, E., Bossinger, O. & Von Mikecz, A. 2009. In Caenorhabditis elegans Nanoparticle-Bio-Interactions Become Transparent: Silica-Nanoparticles Induce Reproductive Senescence. PLOS ONE, 4, e6622.

Ramirez, E., Monteagudo, J. L., Garcia-Gracia, M. & Delgado, J. M. R. 1983. Oviposition and development of Drosophila modified by magnetic fields. Bioelectromagnetics, 4, 315–326.

Shaham, S. W. 2006. Methods in Cell Biology, WormBook. In: Viktor Ambros (ed.) WormBook

Shaw, J., Boyd, A., House, M., Woodward, R., Mathes, F., Cowin, G., Saunders, M. & Baer, B. 2015. Magnetic particle-mediated magnetoreception. Journal of the Royal Society Interface, 12, 0499.

Shcherbakov, D., Winklhofer, M., Petersen, N., Steidle, J., Hilbig, R. & Blum, M. Magnetosensation in zebrafish. Current Biology, 15, R161–R162.

Shtonda, B. B. & Avery, L. 2006. Dietary choice behavior in Caenorhabditis elegans. The Journal of experimental biology, 209, 89–102.

Sulston, J. E., Hodgkin, J. 1988. Methods. In: Wb (ed.) The Nematode Caenorhabditis elegans. Cold Spring Harbor, NY: Cold Spring Harbor Laboratory Press.

Teodori, L., Grabarek, J., Smolewski, P., Ghibelli, L., Bergamaschi, A., De Nicola, M. & Darzynkiewicz, Z. 2002. Exposure of cells to static magnetic field accelerates loss of integrity of plasma membrane during apoptosis. Cytometry, 49, 113–118.

Ueno, S., Lövsund, P. & Öberg, P. Å. 1986. Effects of alternating magnetic fields and low-frequency electric currents on human skin blood flow. Medical and Biological Engineering and Computing, 24, 57–61.

Vidal-Gadea, A., Bainbridge, C., Clites, B., Palacios, B. E., Bakhtiari, L., Gordon, V. & Pierce-Shimomura, J. 2018. Response to comment on “Magnetosensitive neurons mediate geomagnetic orientation in Caenorhabditis elegans”. eLife, 7, e31414.

Vidal-Gadea, A., Ward, K., Beron, C., Ghorashian, N., Gokce, S., Russell, J., Truong, N., Parikh, A., Gadea, O., Ben-Yakar, A., Pierce-Shimomura, J. 2015. Magnetosensitive neurons mediate geomagnetic orientation in Caenorhabditis elegans. Elife, 4.

Wang, L., Du, H., Guo, X., Wang, X., Wang, M., Wang, Y., Wang, M., Chen, S., Wu, L. & Xu, A. 2015. Developmental abnormality induced by strong static magnetic field in Caenorhabditis elegans. Bioelectromagnetics, 36, 178–189.

Wang, L., Wang, M., Du, H., Liu, Y. & Xu, A. 2017. Lipid Metabolism was Interfered by Phosphatidylcholine-Coated Magnetic Nanoparticles in C. elegans Exposed to 0.5 T Static Magnetic Field. Journal of Nanoscience and Nanotechnology, 17, 3172–3180.

Ward, A., Liu, J., Feng, Z. & Shawn Xu, X. Z. 2008. Light-sensitive neurons and channels mediate phototaxis in C. elegans. Nature neuroscience, 11, 916–922.

Wu, Q., Li, Y., Tang, M. & Wang, D. 2012. Evaluation of Environmental Safety Concentrations of DMSA Coated Fe(2)O(3)-NPs Using Different Assay Systems in Nematode Caenorhabditis elegans. PLoS ONE, 7, e43729.

Yang, Y.-F., Lin, Y.-J. & Liao, C.-M. 2017. Toxicity-based toxicokinetic/toxicodynamic assessment of bioaccumulation and nanotoxicity of zerovalent iron nanoparticles in Caenorhabditis elegans. International Journal of Nanomedicine, 12, 4607–4621.

Yemini, E., Jucikas, T., Grundy, L. J., Brown, A. E. X. & Schafer, W. R. 2013. A database of C. elegans behavioral phenotypes. Nature methods, 10, 877–879.

Zablotskii, V., Polyakova, T., Lunov, O. & Dejneka, A. 2016. How a High-Gradient Magnetic Field Could Affect Cell Life. Scientific Reports, 6, 37407.

Zablotskii, V., Syrovets, T., Schmidt, Z. W., Dejneka, A. & Simmet, T. 2014. Modulation of monocytic leukemia cell function and survival by high gradient magnetic fields and mathematical modeling studies. Biomaterials, 35, 3164–3171.

## References

Fonnum, G., Johansson, C., Molteberg, A., Mørup, S. & Aksnes, E. 2005. Characterisation of Dynabeads^®^ by magnetization measurements and Mössbauer spectroscopy. Journal of Magnetism and Magnetic Materials, 293, 41–47.

Shevkoplyas, S. S., Siegel, A. C., Westervelt, R. M., Prentiss, M. G. & Whitesides, G. M. 2007. The force acting on a superparamagnetic bead due to an applied magnetic field. Lab Chip, 7, 1294–302.

